# Controlled Blood-Brain Barrier Modulation by a High-Affinity Claudin-5 Peptide Binder

**DOI:** 10.64898/2026.09.12.751133

**Authors:** Alessandro Berselli, Martina Trevisani, Giulio Alberini, Aldo Pastore, Andrea Di Fonzo, Andrea Armirotti, Valentina Castagnola, Luca Maragliano, Fabio Benfenati

**Affiliations:** Center for Synaptic Neuroscience and Technology, Istituto Italiano di Tecnologia, Largo Rosanna Benzi, 10, 16132 Genova, Italy; Fondazione Pisana per la Scienza ETS, Via Ferruccio Giovannini 13, 56017 San Giuliano Terme, Pisa, Italy; Analytical Chemistry Lab, Istituto Italiano di Tecnologia, Via Morego 30, 16163 Genova, Italy; Department of Life and Environmental Sciences, Polytechnic University of Marche, Via Brecce Bianche, 60131, Ancona, Italy; AOM IRCCS Ospedale Policlinico San Martino, Largo Rosanna Benzi, 10, 16132 Genova, Italy

**Author notes:** Corresponding authors: Valentina Castagnola Luca Maragliano Fabio Benfenati. These authors have contributed equally (Alessandro Berselli, Martina Trevisani, Valentina Castagnola, Luca Maragliano, Fabio Benfenati). Alessandro Berselli, Atomistic Simulations, Center for Human Technologies, Istituto Italiano di Tecnologia, 16156 Genova, Italy. Martina Trevisani, Translational Cancer Medicine Program, 00014 University of Helsinki, Finland.

**Keywords:** Tight junctions, paracellular permeability, peptidomimetics, generative modeling, proteomics

## Abstract

The blood-brain barrier (BBB) is a specialized interface that tightly regulates the exchange of molecules between the bloodstream and the brain. Its barrier function relies on a monolayer of brain endothelial cells sealed by tight junctions (TJs) that restrict paracellular flux through claudin-5 (CLDN5) multimeric complexes. To improve the delivery of nutrients and drugs to the brain, CLDN5-competitive peptides are promising carriers for creating size-controlled, temporary openings in the paracellular pathway. Here, we combine generative protein design and atomistic simulations to design ST9, a peptide with high nanomolar affinity for CLDN5. Compared with f1-C5C2, a CLDN5-binding peptide that we previously reported, ST9 induces a rapid, transient, size-controlled, and fully reversible increase in paracellular permeability, without altering CLDN5 expression or the proteomic profile of brain endothelial cells, indicating distinct mechanisms of TJ destabilization. This work provides a promising approach for developing next-generation BBB-opening agents to effectively treat neurological diseases.

**Highlights:**

- A new peptide design strategy, using generative modeling based on RFdiffusion, identified *de novo* a lead CLDN5-binding peptide;
- The new ST9 peptide displayed a high-nanomolar affinity for CLDN5, high solubility, and absent toxicity;
- *ST9* induces a rapid, transient, size-controlled, and fully reversible increase in paracellular permeability;
- ST9-induced paracellular permeabilization was not associated with CLDN5 degradation, altered proteomic profile, or endothelial apoptosis.

## INTRODUCTION

The treatment of severe brain diseases, including neurological disorders, brain cancer, and genetic conditions associated with impaired delivery of essential molecules to the brain, remains one of the major biomedical challenges of our time. A notable example is glucose transporter-1 deficiency syndrome (GLUT1DS), a rare neurodevelopmental disorder caused by a loss-of-function mutation in the SLC2A1 gene encoding the Glut1 transporter in brain endothelial cells, which prevents an appropriate supply of glucose to the brain^1–3^. Despite a detailed understanding of the molecular and physiological mechanisms underlying GLUT1DS, current treatment options remain limited and predominantly rely on ketogenic dietary therapy, which provides alternative energy substrates to the brain^2^.

The main limitation to disease-modifying treatments for many of these pathologies is the presence of the blood–brain barrier (BBB), a highly specialized vascular interface that tightly regulates molecular exchange between the bloodstream and the central nervous system (CNS)^4,5^. The BBB is formed by a monolayer of brain endothelial cells that control transcellular transport *via* specific transporters and ion channels and are laterally sealed by filamentous protein complexes known as tight junctions (TJs), which effectively prevent the nonspecific diffusion of substances through the paracellular spaces. However, its stringent selectivity also severely restricts the passage of most therapeutic agents, posing a critical obstacle to effective treatments^6^. This challenge has spurred extensive efforts to develop innovative strategies to enable brain drug delivery by transiently modulating paracellular permeability. Passive diffusion of solutes through the paracellular pathway is regulated by multimeric complexes of claudin (CLDN) proteins, which constitute the structural backbone of TJ strands^7,8^. In the brain microvasculature, the predominant isoform is claudin-5 (CLDN5), which forms multimeric assemblies that establish a largely nonselective barrier to ions and small solutes^9–13^, thereby playing a central role in maintaining the BBB paracellular impermeability. This functional importance positions CLDN5 as a key molecular target for strategies aimed at transiently and selectively increasing BBB permeability^14–20^.

Structurally, CLDN proteins are folded in a four-helix bundle comprising four transmembrane helices (TM1-4), linked to each other by two extracellular loops (ECL1-2) and one intracellular loop (ICL). In the paracellular space, ECL1 and ECL2 arrange into a five-stranded *β*sheet and are essential for the proliferation of the TJ strands through formation of intracellular (*cis*-) and intercellular (*trans*-) CLDN-CLDN interactions (**Figure S1A**)^21,22^. While no experimental data have yet shed light on the structure of CLDN-based TJ assemblies, a *consensus*, largely based on computational modeling studies, proposes that CLDN5 aggregates form porous scaffolds that constitute the sole pathways for paracellular flux, yet remain impermeable to almost all ions and molecules (**Figure S1B**)^9–11,23–30^. Among the various strategies explored to increase BBB permeability^14,15^, peptides have emerged as promising carriers for delivering therapeutic agents to the CNS^14,17,20,31^. By targeting CLDN ECLs within the paracellular space, these molecules can loosen the CLDN-CLDN interactions, enabling the temporary delivery of drugs and nutrients to the neuronal tissues (**Figure S1C**). In our previous work, we designed a 14-mer peptidomimetic derived from the CLDN5 ECL domain, named f1-C5C2, which binds CLDN5 proteins with a dissociation constant (K_d_) of 68 µM and increases junctional permeability *in vitro*. While treatment with f1-C5C2 successfully increased BBB permeability, it triggered significant endocytosis of CLDN5-peptide complexes in brain endothelial cells, altering CLDN5 localization at cell-cell contacts and exacerbating dysregulation of the TJ strands.

Here, we introduced a new peptide design strategy, moving from structure-based selection to *de novo* sequence design. Over the past few years, machine learning (ML) has become a central tool in protein science, driven by advances in deep learning–based structure prediction and the growing availability of high-quality structural data. In this context, generative methods based on denoising diffusion models have emerged as a powerful class of approaches for designing novel protein structures. Among these, a computational workflow was recently introduced to design protein-binding sequences directly from target structural information. This strategy combines complementary tools to generate, optimize, and validate novel binders with high experimental success rates^33–36^. At its core is RFdiffusion^33^, a diffusion model derived from RoseTTAFold^37^ (RF) that generates novel protein backbones conditioned on predefined geometries or binding interfaces. The resulting backbones are assigned amino acid sequences using ProteinMPNN, and subsequently refined and evaluated with AlphaFold^35^ (AF2), which provides structural confidence metrics for *in silico* assessment^38^.

In this work, we applied this protocol to design *de novo* potential binders targeting CLDN5. A library of ∼4,000 sequences, ranging from 5 to 15 amino acids in length, was generated and rigorously filtered using computational criteria, including AF2 scoring, water solubility, toxicity prediction, and CLDN5-binding affinity assessed through molecular dynamics (MD) simulations and free energy (FE) calculations. From this screening process, the top candidate, ST9 (sequence SSTWFSAET), was identified and experimentally validated. ST9 exhibits a binding affinity for CLDN5 that is two orders of magnitude higher than that of the previously reported peptide f1-C5C2^16^, and induces a rapid and fully reversible increase in paracellular permeability without CLDN5 endocytosis. The distinct features of the BBB paracellular opening by the two peptides uncover different mechanisms of TJ destabilization that depend on the CLDN5 domain to which they dock. This work establishes a general strategy for targeting claudin proteins and, more generally, other therapeutic targets, and provides a promising route toward the development of next-generation BBB-opening agents for effective therapy of neurological diseases.

## RESULTS

### Peptide design and ranking

We employed the RFDiffusion/ProteinMPNN/AF2 protocol to generate 4,000 sequences, ranging from 5 to 15 amino acids in length and designed to target the CLDN5 ECL1 and ECL2 domains. Based on our previous work on peptide-CLDN5 interactions^16^, we specified Val70 and Val154 as hotspot residues to provide limited conditioning for RFdiffusion. Among the structural confidence metrics produced by AF2 (**Figure S1**), the predicted Alignment Error between interacting chains (*pAE_interaction*) provides a reliable estimate of the accuracy of the modeled binding interface^38^. For the designed sequences, the *pAE_interaction* values were mainly concentrated between 22 and 27 Å (**Figure S1A**), with only a small subset of candidates showing values below 20 Å. As a reference, we calculated the corresponding *pAE_interaction* score for f1-C5C2, the CLDN5-binding peptidomimetic developed and characterized in our previous work^16^. This peptide yielded a value of 23.841 Å, consistent with µM binding affinity benchmarks described in Refs ^33,34^ and the dissociation constant experimentally measured in our previous study^16^ (K_d_ = 68.26 μM).

Based on these considerations, we applied a conservative cutoff of 13 Å to identify the most promising candidates for further investigation. Three peptides, named GP9, SA14, and ST9, fulfilled this criterion and were hence selected for toxicity and aqueous solubility assessment. The structural and confidence descriptors for these peptides, together with those of f1-C5C2, are summarized in **Table 1**.

**Table 1.**
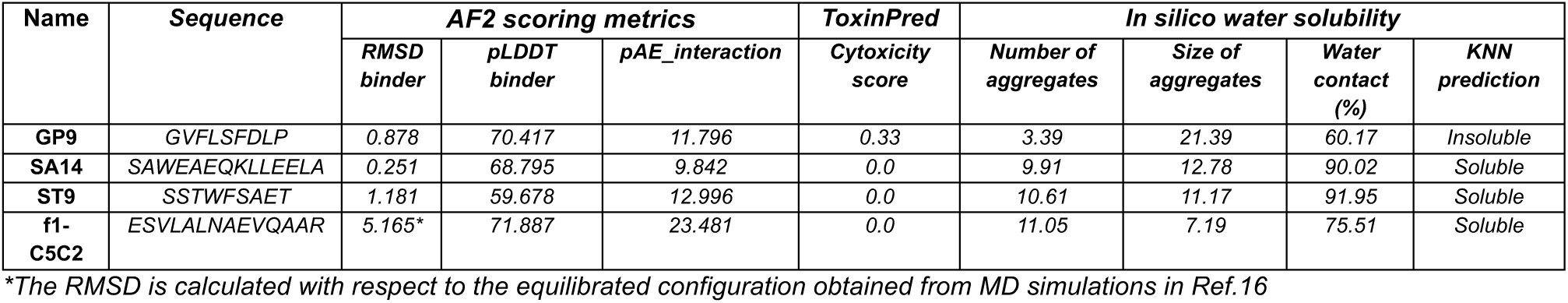
Structural and physicochemical metrics of the designed peptides and f1-C5C2. . AF2 interaction scores, predicted cytotoxicity (ToxinPred), and *in silico* water solubility descriptors are reported.

### Computational assessment of physicochemical properties

Before analyzing CLDN5 binding, we screened the highest-ranked peptides for solubility and predicted cytotoxicity to exclude biologically unsuitable candidates. ToxinPred^39^ classified SA14 and ST9 as non-toxic, while GP9 displayed a cytotoxicity score of 0.33, close to the toxicity threshold of 0.38, suggesting a possible borderline cytotoxic profile.

To predict the solubility of the designed peptides, we used a procedure, optimized in our previous work^16^, tailored to reproduce experimental conditions. As detailed in the **Supplementary Computational Methods** section, the protocol combines MD simulations of collections of peptides with supervised classification (**Figure S3**). A K-nearest neighbors (KNN) classifier is trained on an external set of 16 peptides of known hydrophilicity and used to classify candidates based on MD-derived descriptors: the average size of the largest aggregate, the number of aggregates, and the fraction of water contacts. Simulations (**Figure S3A,B**) revealed that SA14 and ST9 peptides remained significantly dispersed in water at the end of the trajectories, forming ∼10 distinct aggregates with a maximal size of 11-12 molecules (**Figure S3C**). Moreover, the hydrated surface at the end of the simulation was ∼90% of the initial value for both oligomers. In contrast, GP9 formed massive, poorly dispersed aggregates, with only ∼3 aggregates after 100 ns, each with a maximum of ∼22 oligomers. Consistently, the solvent-exposed surface of the peptides at the end of the simulation was only 64% of the initial value. KNN classification (**Figure S3D**) showed a clear distinction between SA14 and ST9, which clustered with water-soluble peptides, and GP9, which was predicted to be insoluble. These results, combined with the potential toxicity evaluation, led us to exclude GP9 for further investigation of the CLDN5-binding affinity.

### Simulations of CLDN5-peptide complexes

The peptides ST9 and SA14 were selected to evaluate their binding affinity to CLDN5. The initial configurations of the protein–peptide complexes were obtained from the AF2-generated refined structural models. Notably, the two peptides exhibit distinct structural organizations and binding modes. ST9 remains largely unfolded and establishes extensive contacts along the entire β4 strand of CLDN5, partially rearranging to act as an extension of the protein’s β-sheet domain (**Figure 1**). In contrast, SA14 adopts a fully folded conformation, forming a long α-helix that primarily interacts with the *β*-turn motifs of the CLDN5 ECL1 (**Figure S4**).

**Figure 1.**
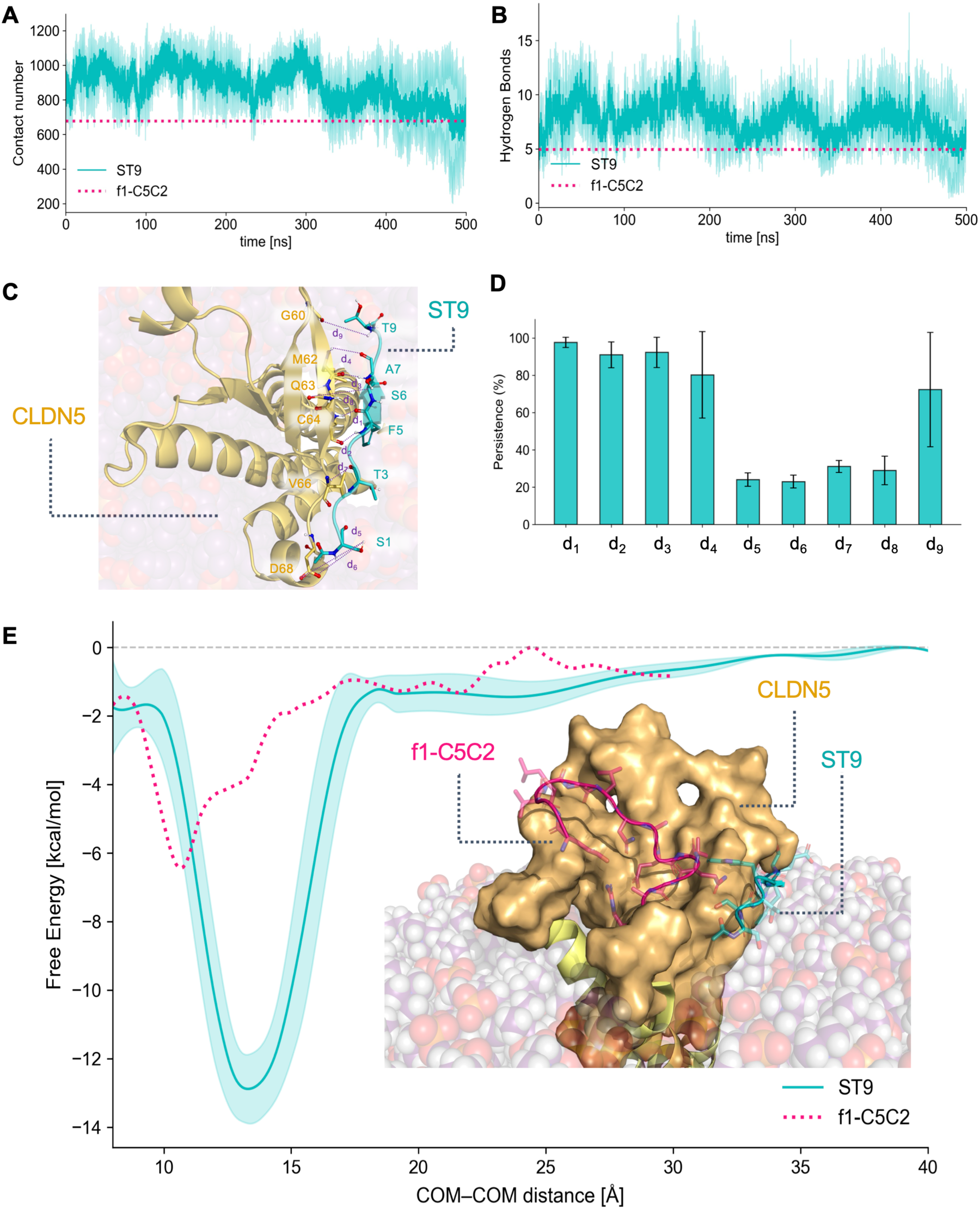
Assessment of CLDN5-ST9 binding affinity via MD simulations and free energy calculations. Time evolution of the number of atom-atom contacts (**A**) and HBs (**B**) between CLDN5 and ST9 during MD simulations. The magenta dotted lines indicate the mean values obtained for f1-C5C2 in Ref.*16*. (**C**) Interaction interface between the CLDN5 ECL domain (yellow) and ST9 (cyan). The most persistent HBs detected during MD simulations are indicated with purple dotted lines. Oxygen, nitrogen, and hydrogen atoms involved in the interaction interface are reported as spheres and colored in red, blue, and white, respectively. (**D**) Persistence of the HBs between CLDN5 and ST9 along MD trajectories, expressed in percent of the simulation time. Only interactions with persistence > 20% are shown. (**E**) Free-energy profile for the unbinding of the CLDN5-ST9 complex. The FE is calculated as a function of the distance between the centers of mass (COMs) of the CLDN5 ECL domain and ST9, and compared to the profile obtained for f1-C5C2 (magenta dotted line, Ref.*16*). The results and associated errors for each analysis are reported as the mean and standard deviation over three independent simulation replicas.

Below, we report the results for ST9, which maintained a stable binding mode throughout MD simulations, and compare them with those for f1-C5C2 reported in Ref.16. Results for SA14, which showed less stable binding, are provided in the **Supplementary Results** section.

First, we monitored the total number of atom-atom contacts between the CLDN5 ECL domain and the peptides (**Figure 1A**). ST9 maintained a stable number of contacts (about 900) for the first 300 ns in all replicas, followed by a decrease at the end of some trajectories. Notably, despite the shorter sequence length, ST9 formed a number of contacts comparable to or higher than the average observed for f1-C5C2 (magenta dashed line), suggesting a more extensive engagement of the new peptide with the CLDN5 surface.

Hydrogen bond (HB) analysis showed that ST9 formed a stable network with CLDN5, maintaining ∼8 HBs on average throughout the simulation, higher than the number observed for f1-C5C2 (**Figure 1B**). ST9 established persistent interactions with CLDN5 along the entire sequence length (**Figure 1C and Table S1**). The HBs formed between the backbones of Phe5, Ser6, and Ala7 in ST9 and those of 5464, Gln63, and Met62 in CLDN5 (named d_1_–d_4_) were maintained for > 80% of the total simulation time. Additional stabilizing interactions involved the side chains of Ser1 and Asp68 (d_5_– d_6_), the backbones of Thr3 and Val66 (d_7_), the side chains of Ser6 and Gln63 (d_8_), and the backbones of Thr9 and Gly60 (d_9_).

Despite the longer sequence, the number of contacts between the SA14 peptide and CLDN5 during MD simulations was significantly lower (∼300), and peptide dissociation was observed in two of the three replicas (**Figure S4A**). Moreover, SA14 maintained < 5 HBs for the entire duration (**Figure S4B**), with only two interactions with an average persistence exceeding 20% of total simulation time, involving the backbones of Ala2 and Ala43 (d_10_) and of Trp3 and Val56 (d_11_). These interactions showed large uncertainties, reflecting the complex’s variable stability across simulation replicas (**Figure S4C,D**).

To quantitatively assess the stability of CLDN5-ST9 binding, we performed FE calculations using Temperature-Accelerated Molecular Dynamics (TAMD)^40,41^. Consistent with our previous work^16^, we selected the distance between the centers of mass (COMs) of the CLDN5 ECL domain and the peptide as a collective variable (CV). The FE was obtained by integrating the mean forces acting on the CV over TAMD trajectory^40,42^. The resulting FE profile (**Figure 1E**) exhibits a deep and relatively sharp minimum of approximately 13 kcal/mol, twice as deep as that obtained for f1-C5C2^16^, at a COM–COM distance of ∼12.7 Å. Hence, the computational results presented here indicate that ST9 is predicted to have a substantially higher binding affinity for CLDN5 than f1-C5C2, supporting its prioritization as a promising candidate for further *in vitro* evaluation.

### Biological effects of ST9 exposure in an in vitro BBB model

Due to the wide availability of murine *in vitro* and *in vivo* models of BBB, we first assessed whether the results of the computational analysis on the human CLDN5 (hCLDN5) protomer as a receptor model^9–11,16^, also apply to the murine CLDN5 isoform (mCLDN5). Notably, the human (UNIPROT: O00501) and murine (UNIPROT: O54942) CLDN5 orthologs share 92% global identity, which increases to >95% within the ECL domains. When the AF2 refinement and scoring of ST9 were repeated with mCLDN5, a *pAE_interaction* of 10.091 was obtained, slightly better than with hCLDN5 and indicative of a strong binding affinity. For this reason, mCLDN5 and mouse bEnd.3 brain endothelial cells were used in the following experimental work.

The binding affinity of ST9 was experimentally validated using microscale thermophoresis (MST), following the same experimental design as in our previous study^16^. The binding isotherms of ST9 to CLDN5 (cyan line) as a function of peptide concentration were markedly left-shifted relative to those of f1-C5C2 (dotted pink line; **Figure 2A**). The obtained nanomolar-scale dissociation constant (K_d_ = 726 nM) was about two orders of magnitude lower than that of f1-C5C2 (68 µM^16^), confirming the enhanced binding affinity of ST9 for CLDN5 predicted by the computational analysis.

**Figure 2.**
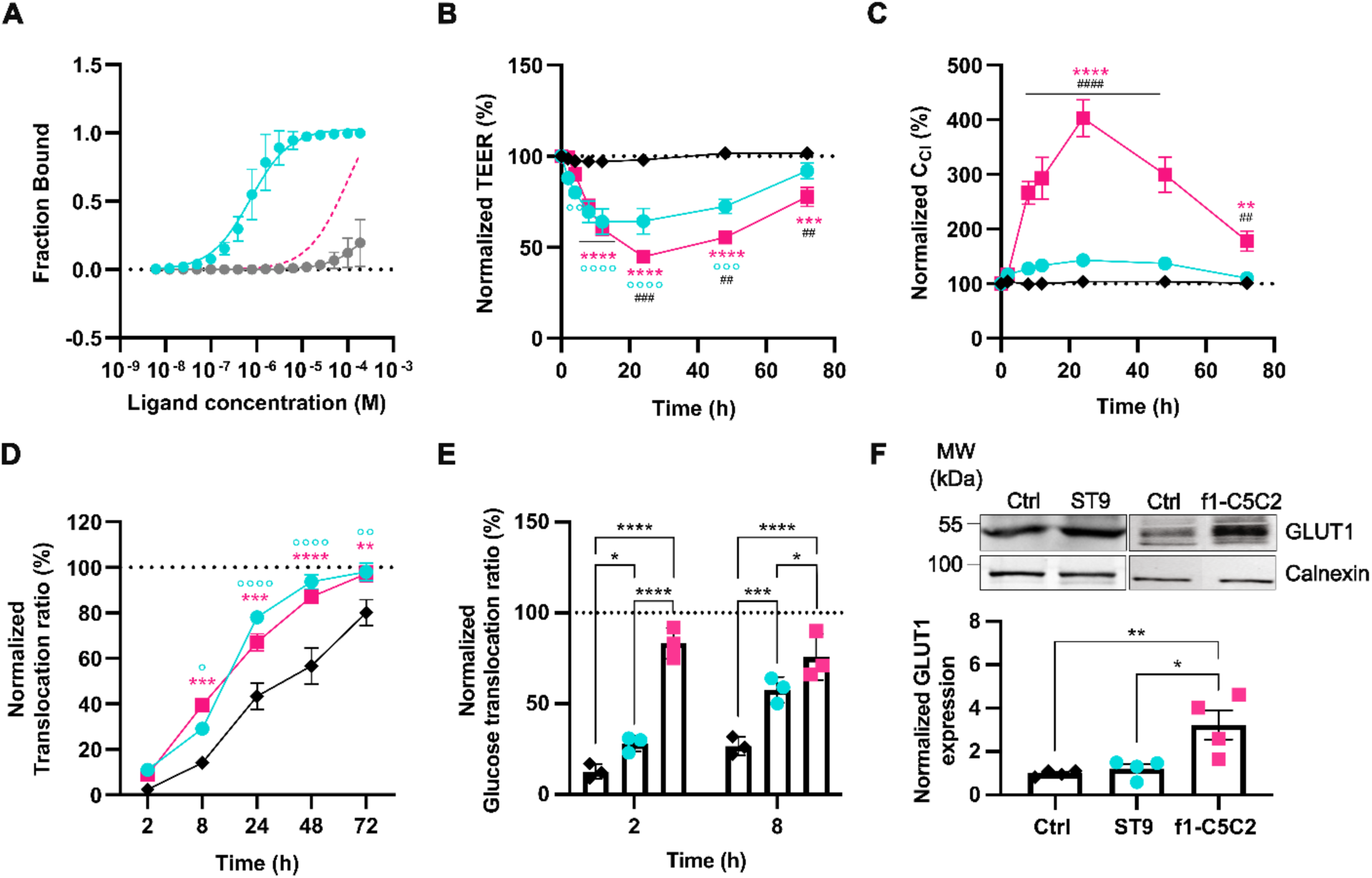
Binding and biological effects of ST9 exposure in comparison with f1-C5C2. (**A**) MST binding curves for ST9 (cyan circles) are shown together with a negative control sample (grey circles, recombinant tGFP spotted in the cell lysate in the absence of transfected mCLDN5) and f1-C5C2 binding curve fitting (pink line, adapted from Ref.*16*). ST9 curve fitting yielded a K_d_ value of 726 nM for ST9. (**B-C**) TEER values (**B**) and C_Cl_ (**C**), presented as percentages of baseline values, were measured in bEnd.3 monolayers treated with ST9 (250 µM, cyan circles) and f1-C5C2 (25 µM, pink squares) for up to 72 hours. The vehicle was used as a control condition (black circles). (**D**) Normalized translocation ratio for FD4 across bEnd.3 cell layers exposed to ST9 (250 µM, cyan circles) and f1-C5C2 (25 µM, pink squares) or vehicle (black). The translocation ratio (expressed as a percentage of the translocation across a Transwell membrane without cells) was calculated as the ratio of dextran mass in the basolateral side at the sampling time point to the apical side at t = 0. Data are shown as means ± SEM (n = 3). *: f1-C5C2 vs Ctrl; °: ST9 vs Ctrl; #: ST9 vs f1-C5C2. *p < 0.05, **p < 0.01, ***p < 0.001, ****p < 0.0001 two-way repeated measures analysis of variance (ANOVA)/Tukey’s tests. (**E**) Normalized translocation ratio for D-Glucose across bEnd.3 cell layers pretreated with ST9 (250 µM, cyan circles), f1-C5C2 (25 µM, pink squares), or vehicle (black) for 24 h. Data are shown as means ± SEM (n = 4). **p < 0.01, ***p < 0.001, ****p < 0.0001 two-way repeated measures analysis of variance (ANOVA)/Tukey’s tests. (**F**) Top: Representative Western blots of GLUT1 (55 kDa) and calnexin (100 kDa) in bEnd.3 cells exposed for 24 hours to either vehicle (Ctrl, black), ST9 (250 µM, cyan circles), or f1-C5C2 (25 µM, pink squares). Bottom: GLUT1 expression was quantified by densitometric analysis, normalized to calnexin, and expressed relative to Ctrl. Data are shown as means ± SEM; individual dots represent independent replicates. *p < 0.05, **p < 0.01; one-way ANOVA/Tukey’s multiple-comparisons test.

Next, we employed a widely used 2D in vitro model of the BBB with murine bEND.3 brain endothelial cells in a Transwell^®^ system to investigate the biological activity of the ST9 peptide and compare it with the f1-C5C2 peptide^16^. We first carried out a dose-response study of the ST9 peptide by evaluating the trans-endothelial electrical resistance (TEER). We found that, despite the sub-micromolar K_d_ for CLDN5, we had to increase the ST9 concentration above 100 μM to yield a significant decrease in TEER values starting after 4 h of incubation (**Figure S5**). We then compared the effects of ST9 with those previously described for f1-C5C2^16^. The latter peptide could not be used at concentrations above 25 µM due to solubility issues and morphological changes in endothelial cells^16^. In the case of ST9, the TEER values (**Figure 2B**) and the relative cell layer capacitance (C_cl_; **Figure 2C**) followed the kinetics previously observed for f1-C5C2, reaching a minimum after 24 h of continuous incubation. However, ST9 showed a slightly smaller overall decrease in TEER and complete recovery to control values after 72 h. No significant effects were observed with the SA14 peptide, used as a negative control, on MST, TEER, and C_cl_ values (**Figure S6**). Permeability assays reported in **Figure 2D** (for FITC-dextran 4 kDa – FD4) and **Figure S7** (FITC-dextran 40 kDa and 70 kDa – FD40 and FD70) showed very comparable results between the two peptides (see also Ref.16). This size screening indicates that the temporary barrier opening permits the passage of species with a hydrodynamic diameter (D_h_) below 5 nm, while not affecting the translocation of species with D_h_ ≥ 7–10 nm, as observed for FD70 (**Figure S7**).

Finally, to assess the potential translational relevance to GLUT1DS, the effects on glucose translocation were measured 2 and 8 h after a 24-h pretreatment of bEND.3 cells with either ST9 or f1-C5C2, using a D-glucose colorimetric detection kit. ST9 significantly increased glucose translocation at 2 h, although to a lesser extent than f1-C5C2. Notably, the ST9-induced glucose translocation increased markedly at 8 h, approaching the levels observed with f1-C5C2 (**Figure 2E**). However, in the latter case, the increased glucose passage was associated with GLUT1 upregulation, which prevented isolating the paracellular transport contribution from the transcellular one (**Figure 2F)**. In contrast, the significant increase in glucose transport following ST9 administration was solely attributable to enhanced paracellular permeability, as no significant changes in GLUT1 expression were observed by quantitative immunoblot analysis.

To elucidate the downstream mechanistic action following peptide binding to the extracellular portion of CLDN5, we compared the expression of CLDN5 and other relevant TJs proteins (Occludin and *Zonula Occludens*, ZO1) and monitored CLDN5 subcellular localization over time. Data were collected after 24 h of continuous exposure, when the maximum barrier opening (minimum TEER) was observed. In our previous work, we observed that 24 h of continuous exposure to f1-C5C2 resulted in a significant downregulation of TJ protein expression (**Figure 3A-C**, dotted bars). On the contrary, exposure to ST9 did not affect TJs expression, which remained comparable to that in untreated controls. Interestingly, CLDN5 not only maintained stable expression levels but also maintained its correct membrane localization, which is necessary for fully reconstituting the tetrameric assembly (**Figure 3D,E**). This behavior of ST9 suggests that the peptide acts via a different mechanism of action with respect to f1-C5C2, which induced a massive downregulation and cellular internalization of CLDN5^16^.

**Figure 3.**
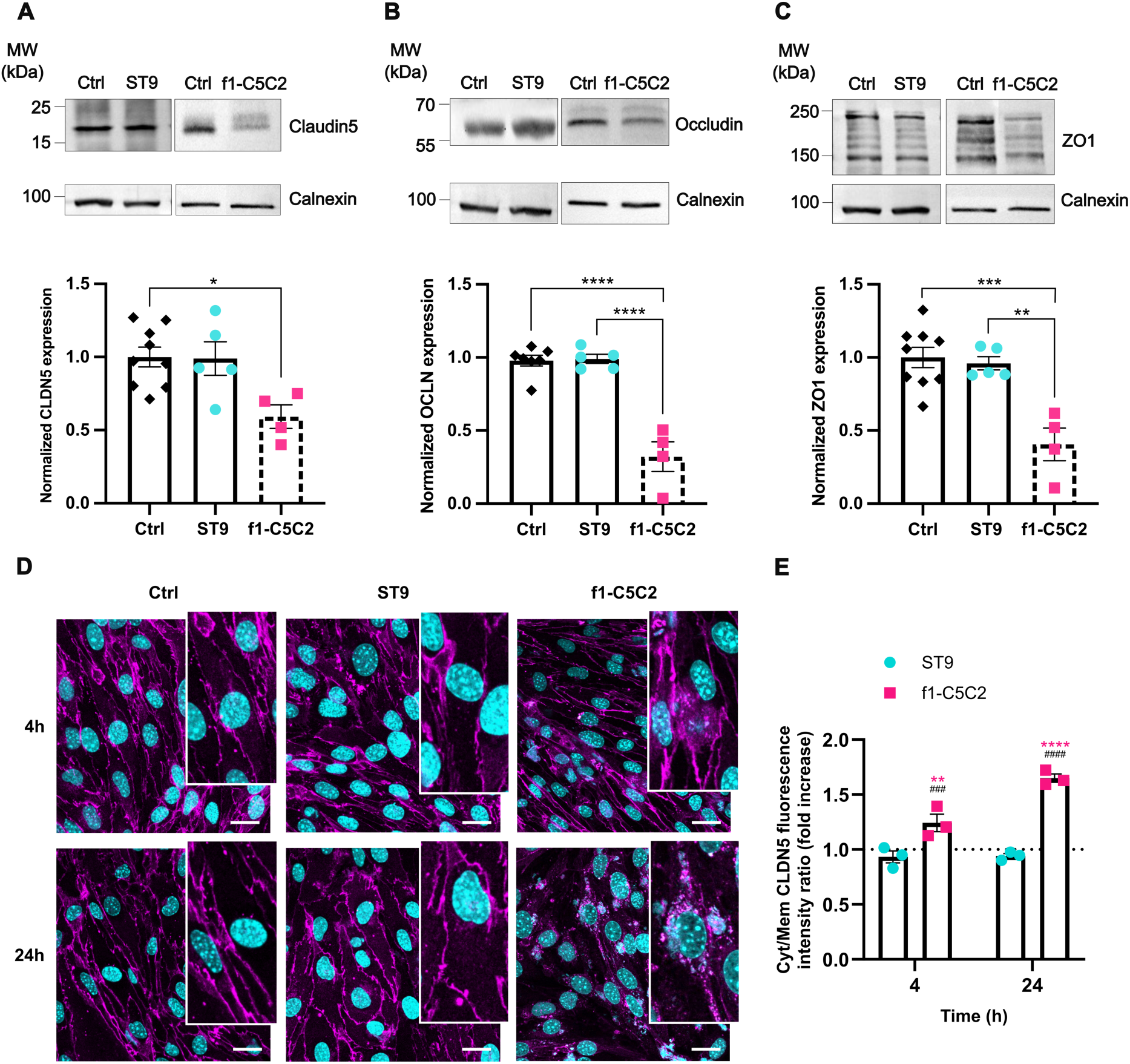
Tight-junction protein expression and intracellular localization upon peptide binding. (**A-C**) *Top:* Representative Western blots stained with CLDN5 [(**A**) 23 kDa], OCLN [(**B**) 55 kDa], ZO1 [(**C**) 220 kDa], and calnexin (100 kDa) for bEnd.3 cells after 24 h of treatment with either vehicle (Ctrl, black), 250 µM ST9 (cyan), or 25 µM f1-C5C2. *Bottom:* Quantification of protein expression normalized to the respective controls. Data are means ± SEM (n = 5). (**D**) Representative confocal images of CLDN5 immunostaining at 4 and 24 h for bEnd.3 cells exposed to vehicle (Ctrl), 250 µM ST9 or 25 µM f1-C5C2. Scale bars, 25 μm. (**E**) Quantification of CLDN5 intracellular redistribution based on image analysis. For each time point, the cytoplasmic-to-plasma membrane CLDN5 fluorescence intensity ratio was calculated for each cell and normalized to the ratio measured in matched controls. Data are means ± SEM (n = 3 independent experiments in triplicate; 5 fields per replicate). *: f1-C5C2 vs Ctrl; #: ST9 vs f1-C5C2. **p < 0.01, ***p < 0.001, ****p < 0.0001; two-way repeated measures ANOVA)/Tukey’s tests.

Viability analysis showed no PI-positive, necrotic cells with either peptide at the respective concentrations (**Figure 4A**). However, f1-C5C2 at 25 µM induced apoptosis in ∼15% of bEnd.3 cells, preventing the use of higher concentrations. This observation might explain the significantly higher values recorded for C_cl_ (**Figure 2C**) and underlie the observed increased expression of GLUT1 (see **Figure 2F**) observed after incubation with f1-C5C2. In contrast, ST9 exhibited high biocompatibility, enabling the use of higher concentrations without effects on cell viability, and its effect on TEER was fully reversible, once again suggesting a distinct mechanism of action.

**Figure 4.**
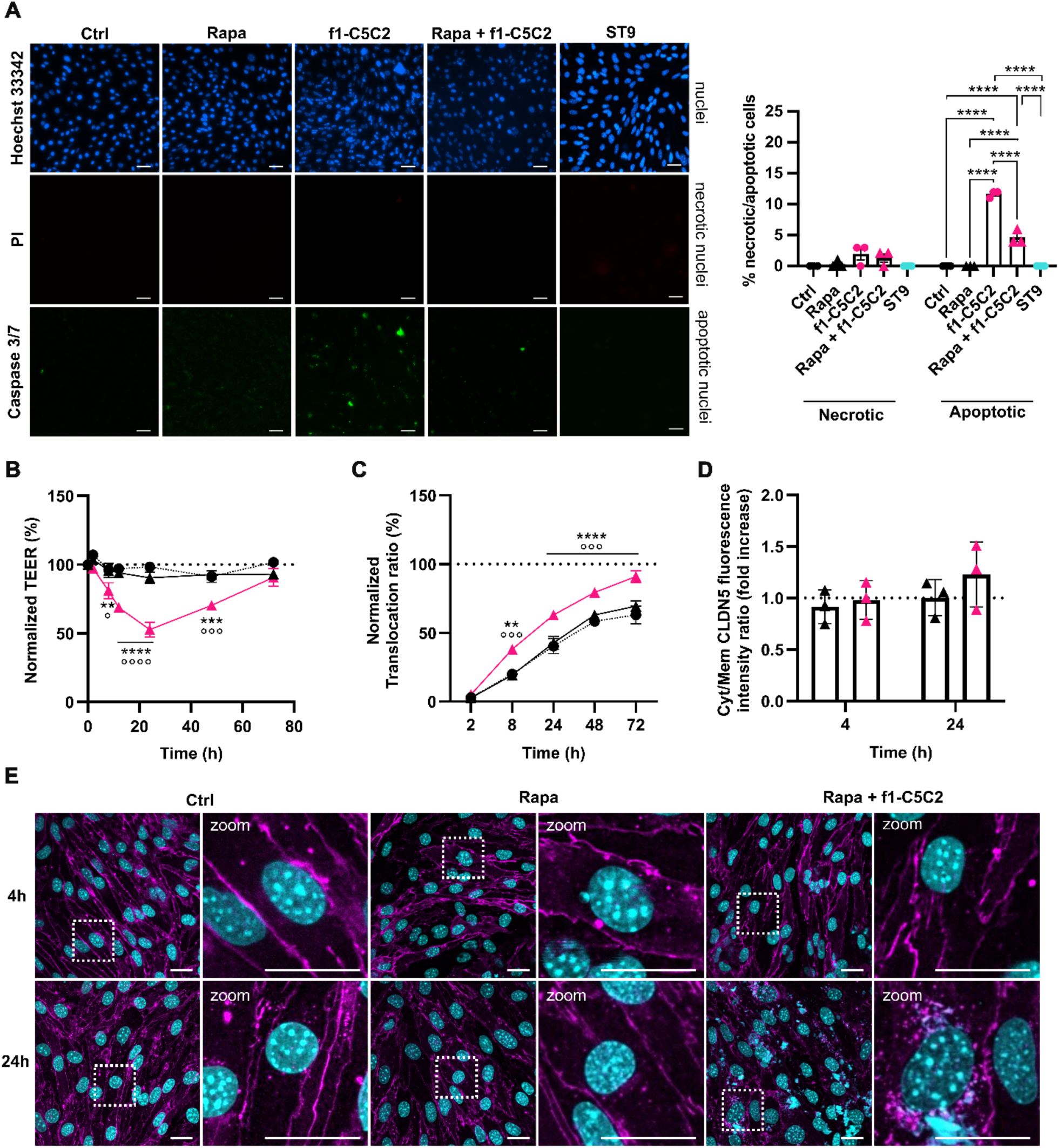
Rapamycin-induced autophagy improves viability and barrier recovery after f1-C5C2 treatment. (**A**) *Left:* Representative fluorescence images at t = 4 h for bEnd.3 cells treated with vehicle (Ctrl), 25 μM f1-C5C2, rapamycin alone, 25 μM f1-C5C2 + rapamycin, or 250 μM ST9. Cells are stained with Hoechst 33342 (nuclear staining), Propidium iodide (PI, necrotic cells), and Caspase 3/7 (apoptotic cells). Scale bar, 50 µm. *Right:* Quantification of the amount of necrotic and apoptotic cells in percent of total cells. Data are means ± SEM (n = 3 independent experiments; five-seven fields per replicate). ****p < 0.0001 by two-way repeated-measures ANOVA/Tukey’s tests *versus* control. (**B,C**) TEER values (**B**) and normalized translocation ratio for FD4 (**C**) measured in bEnd.3 monolayers treated with vehicle (Ctrl, black circles), rapamycin alone (black triangles), or 25 μM f1-C5C2 + rapamycin (pink triangles) for up to 72 h. Data are shown as means ± SEM (n = 3). *: Rapa + f1-C5C2 vs Ctrl; °: Rapa + f1-C5C2 vs Rapa. *p < 0.05, **p < 0.01, ***p < 0.001, ****p < 0.0001 two-way repeated measures ANOVA/Tukey’s tests. (**D**) Quantification of CLDN5 intracellular localization based on image analysis. For each time point, the cytoplasmic-to-plasma membrane CLDN5 fluorescence intensity ratio was calculated per cell and normalized to the ratio measured in matched controls. Data are means ± SEM (n = 3 independent experiments in triplicate; five fields per replicate). (**E**) Representative confocal images of CLDN5 immunostaining in bEnd.3 cells treated with vehicle (Ctrl), Rapamycin, or 25 μM f1-C5C2 + Rapamycin, for 4 and 24 h. Scale bar, 25 μm.

The internalization of CLDN5 (see **Figure 3D,E**) and the degree of apoptosis in endothelial cells treated with f1-C5C2 (**Figure 4A**) are likely attributable to a dysregulation of autophagy, which has been reported to associate with apoptotic cell death, barrier dysfunction, and accelerated cytosolic degradation of CLDN5^44,45^. Based on these considerations, we examined whether inducing autophagy could attenuate the apoptosis observed following exposure to f1-C5C2. Autophagy was induced by treating cells with the mTORC1 inhibitor Rapamycin (Rapa; **Figure 4**) or by starvation for 24 h in DMEM supplemented with 2% FBS (Starv; **Figure S8** and **S9**). Autophagy induction by either protocol reduced f1-C5C2-induced apoptosis to approximately 5% and, accordingly, abolished CLDN5 internalization. However, the treatments only minimally attenuated the f1-C5C2-induced decrease in TEER and the enhanced FD4 permeability.

Proteomic analyses of bEND.3 cells incubated with either f1-C5C2 or ST9 support the hypothesis that binding to distinct sites on CLDN5 activates different downstream pathways in endothelial cells (**Figure 5**). The Volcano plots in **Figure 5** show that, while exposure to ST9 did not significantly alter the proteome compared to untreated cells (**Figure 5A**), exposure to f1-C5C2 resulted in several proteins that were significantly down-or upregulated (**Figure 5B**).

**Figure 5.**
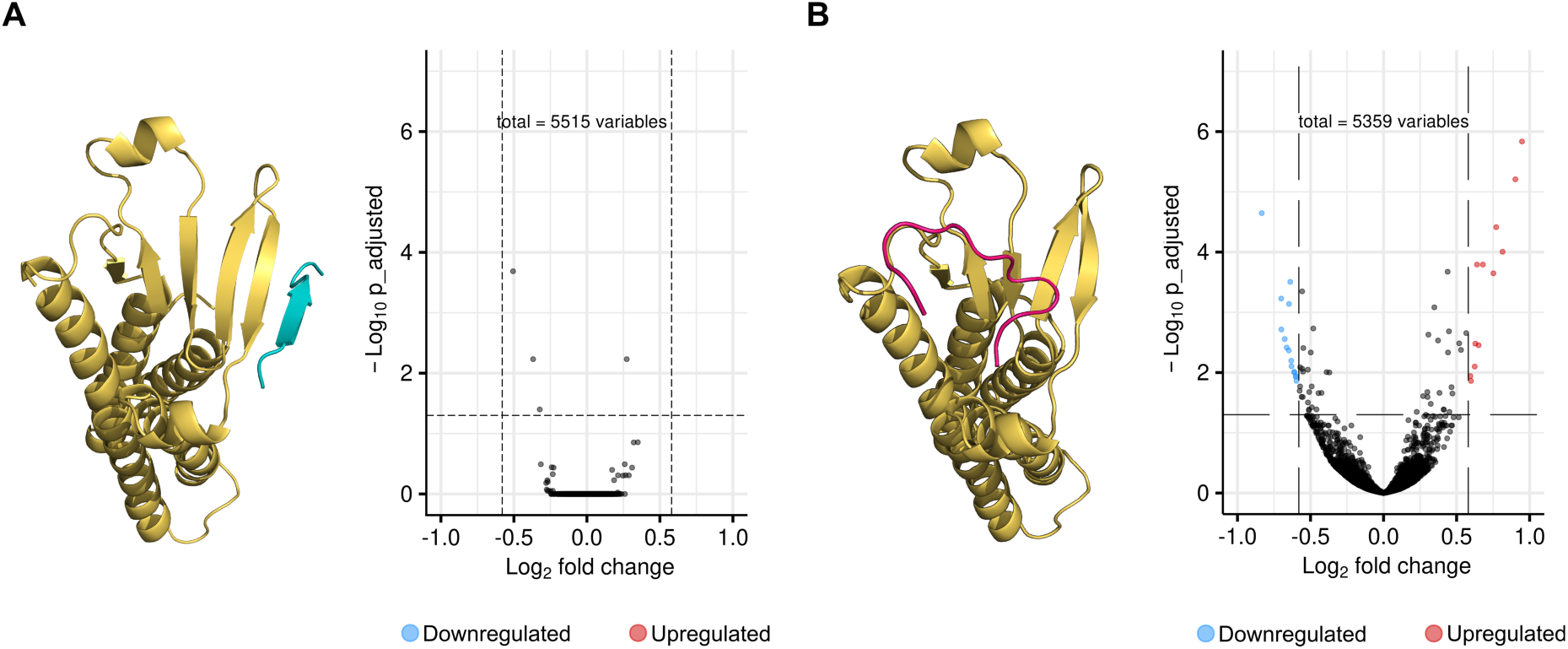
Proteomic responses induced by ST9 and f1-C5C2 upon CLDN5 targeting. (**A, B**) *Left:* Representative binding models of ST9 (**A**, cyan) and f1-C5C2 (**B**, magenta) interacting with the CLDN5 complex (yellow). *Right:* Volcano plots showing differentially regulated proteins following exposure to ST9 (**A**) or f1-C5C2 (**B**). Blue and red dots indicate significantly downregulated and upregulated proteins, respectively, while grey dots represent proteins that did not reach statistical significance. Dashed lines indicate the applied fold-change and adjusted P-value thresholds.

The analysis of the top enriched pathways associated with differentially regulated proteins following exposure to f1-C5C2, focusing on cellular localization–related terms (**Figure 6A**), revealed a prominent interaction network centered on mitochondrial activity, which is closely linked to the observed dysregulation of apoptosis. In addition to caspases, which are prime mediators of apoptosis, the markedly altered network of serine endopeptidases^46^ (serine proteases), observed in the molecular function-related terms (**Figure 6B**), plays critical, complementary roles in promoting, regulating, and executing apoptosis. As a result, vasculature development and morphogenesis (biological process-related terms; **Figure 6C**) are affected, as cell death regulators are closely linked to blood vessel regression and remodeling^47,48^. The study of the effects of peptide incubation over time, shown in **Figure S10** as Venn diagrams of enriched pathways for biological process, cellular component, and molecular function, confirms that the number of enriched pathways increases with exposure time to f1-C5C2. Altogether, the bioinformatics analysis of the whole proteome of brain endothelial cells fully supports a distinct mechanism of action of f1-C5C2 relative to ST9, for which the only affected network pathway in molecular function-related terms is the ligase catalytic activity (**Figure 6B**).

**Figure 6.**
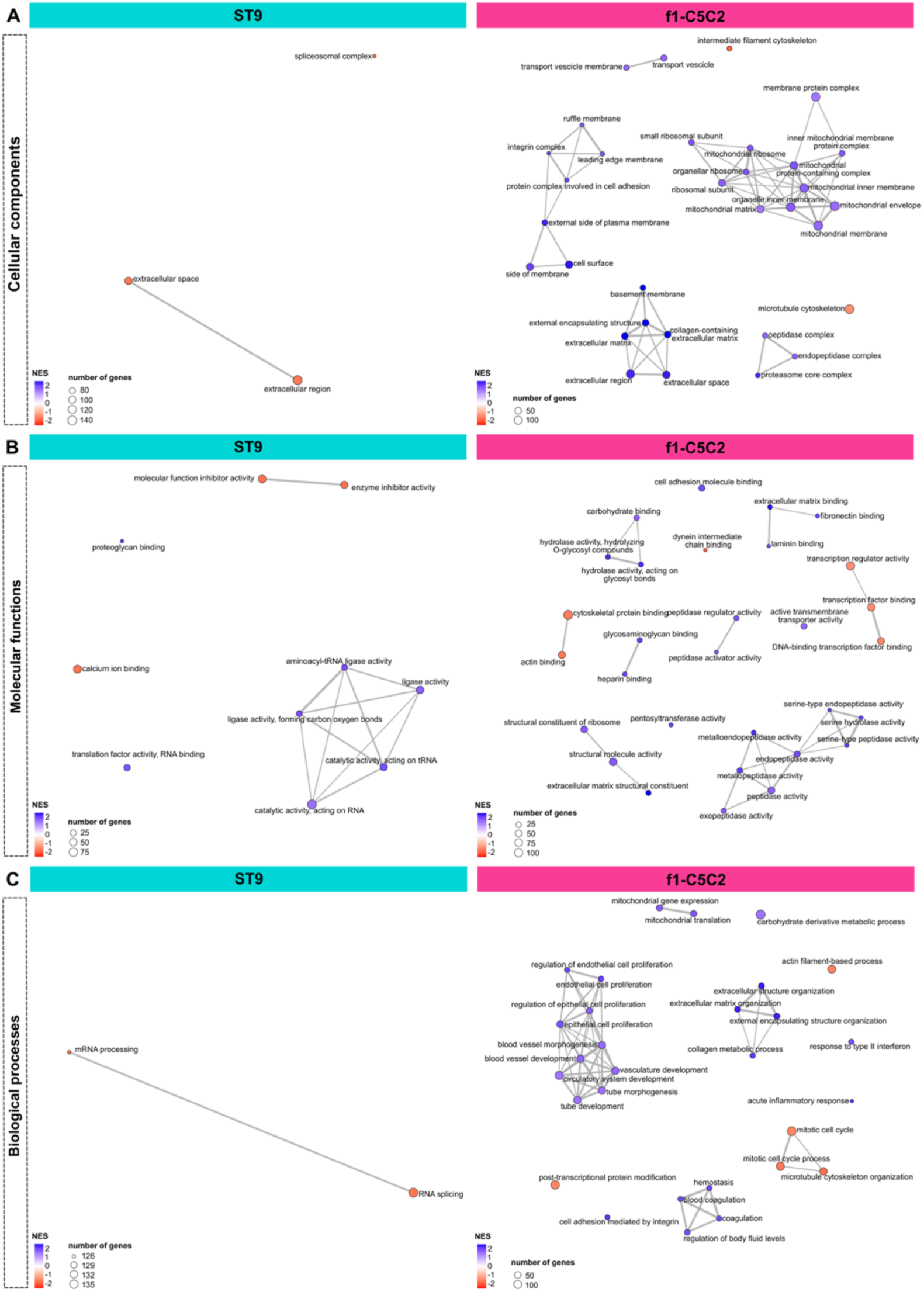
Protein-based functional enrichment analysis on bEND.3 cells after 24 h exposure to ST9 and f1-C5C2, respectively. (**A-C**) Network visualization of the top enriched pathways associated with differentially regulated proteins following exposure to ST9 (cyan, left panels) or f1-C5C2 (magenta, right panels). Enrichment maps are shown for cellular localization-related terms (**A**), molecular function-related terms (**B**), and biological process-related terms (**C**). Nodes represent enriched protein-associated terms, with node size proportional to the number of proteins included in each term and node color indicating the Normalized Enrichment Score (NES). A positive NES (blue) indicates an activated pathway, whereas a negative NES (red) indicates a suppressed pathway. Edges connect terms sharing common proteins.

## DISCUSSION

The controlled delivery of therapeutic agents to the CNS for the treatment of neurological diseases is fundamentally constrained by the BBB. The significant advances made in recent years in the molecular-level understanding of CLDN-based TJ strands^8,11^ have offered unique opportunities for the rational design of peptide binders targeting the intermolecular interactions that underlie the stability of their multimeric architecture^16,17,20^. Peptide-based TJ modulators offer several advantages in this context, including modular and flexible engineering, systemic biodistribution compared to the localized physical approaches such as focused ultrasound-mediated BBB opening^49^, and inherently transient action arising from their short half-life in biological environments^50^. Peptides offer the specificity of large biological molecules, such as antibodies, while retaining the size and synthetic features of small-molecule drugs^51^. Importantly, appropriately designed peptides can enable transient, controlled, and non-toxic modulation of BBB permeability, making them particularly attractive for clinical translation.

Leveraging structural information from multi-pore CLDN5 models, we previously designed the peptidomimetic f1-C5C2, which displayed measurable binding affinity for CLDN5 (K_d_ = 68.26 μM) but induced undesired cellular effects, including CLDN5 internalization and partial apoptosis^16^. Rather than iteratively refining this sequence by further targeting the intermolecular contacts observed in multimeric CLDN5 models, we here revisited our design workflow by adopting state-of-the-art generative models, RFDiffusion^33^ and ProteinMPNN^38^, to produce *de novo* peptide candidates. These deep-learning-powered methods represent a paradigm shift in protein and peptide design^52^, moving from energy-guided structural screening grounded in the assumption that optimal configurations correspond to free energy minima^53^, to data-driven approaches in which the information required to generate viable structures is encoded in the large experimental datasets used to train the models. While the predictive power of these frameworks has been demonstrated across a broad range of protein-ligand scenarios, including antibodies^54^, mini-proteins^33,36^, and linear and cyclic peptides^55,56^, experimental success with peptide binders remains more heterogeneous than for protein scaffolds and is critically dependent on the successive steps of computational filtering and orthogonal validation^33,55,57^.

For this reason, while our preliminary filtering of the 4000 ML-generated sequences promoted 3 candidates for further characterization, a meticulous MD simulation-based analysis was essential to refine the selection: it allowed us to exclude the potentially toxic and insoluble candidate GP9 and, importantly, to identify that the apparently best-performing sequence SA14, based on AF2 scoring metrics alone, does not bind CLDN5 efficiently in the explicit-solvent MD regime. These results reinforce the rationale behind our pipeline, which combines the high-throughput generative power of ML models with the detailed atomistic description of the binding event provided by MD simulations and underscore that AF2 confidence metrics alone are insufficient surrogates for binding affinity in the context of flexible peptide-protein interactions.

Our workflow ultimately yielded ST9, a 9-residue peptide with a K_d_ of 726 nM, 2 orders of magnitude lower than f1-C5C2. Structurally, ST9 establishes a stable, geometrically optimized network of HBs along the entire β4 strand of the CLDN5 ECL1 domain, effectively extending the protein β-sheet via a strand-augmentation mechanism. Interestingly, this binding mode was not anticipated *a priori but* emerged naturally from the generative design process. In contrast, f1-C5C2 interacts primarily with the helical and unstructured regions of the ECL domain, which are intrinsically less geometrically constrained, resulting in a less persistent, lower-affinity binding mode. The quantitative agreement between the computed FE barriers (∼13 kcal/mol for ST9 vs. ∼6 kcal/mol for f1-C5C2) and the experimentally measured K_d_ values further validates the MD/TAMD-based affinity ranking as a predictive tool within our design workflow.

Beyond differences in binding affinity, a crucial result is the fundamentally distinct mechanisms by which ST9 and f1-C5C2 operate in the biological environment. Exposure to f1-C5C2 induced marked downregulation of CLDN5, occludin, and ZO-1, massive redistribution of CLDN5 from the plasma membrane to the cytoplasm, increased GLUT1 expression, and activation of apoptotic pathways, resulting in dysregulation of the whole proteome of the brain endothelial cells.

Another notable difference concerns the effective working concentrations. While f1-C5C2 induced a decrease in TEER at 25 µM (a concentration at the solubility limit) and no further effect was observed at higher concentrations^16^, ST9 (solubility limit >250 µM) produced a measurable perturbation of TEER at concentrations above 100 µM. It is tempting to speculate that f1-C5C2 acts through a more “*catalytic*” mechanism, whereby a limited number of peptide molecules is sufficient to trigger CLDN5 intracellular reorganization. In contrast, ST9 appears to exert a more “*stoichiometric*” effect on the intact CLDN5-CLDN5 interface, requiring saturation of CLDN5 binding sites to produce a macroscopic change in barrier integrity, without triggering secondary amplification mechanisms such as massive CLDN5 recycling and downregulation.

We propose that f1-C5C2 binding to the exposed, unstructured ECL regions (**Figure 7**) triggers internalization and autophagic degradation of CLDN5, similar to the mechanism recently described for the C-terminal fragment of the *Clostridium perfringens* enterotoxin (cCPE) in brain endothelial cells^58^. In the latter case, caveolin-1 mediated the internalization of CLDN5 following cCPE binding. The aberrant cytosolic accumulation triggered autophagic degradation of CLDN5, ultimately increasing BBB permeability. In different barrier settings, such as the intestinal barrier, inhibition of autophagy exacerbates barrier dysfunction characterized by reduced expression of CLDN1 and ZO1^59^. In this framework, the apoptotic response observed upon f1-C5C2 treatment is best understood as a secondary consequence of impaired autophagy rather than a direct cytotoxic effect of the peptide, a hypothesis supported by the partial rescue of cell viability observed upon autophagy activation by rapamycin or starvation. This mechanistic difference is directly reflected in the reversibility and safety profile of the BBB modulation. ST9 administration led to complete TEER recovery within 72 h, with negligible apoptosis at the working concentration of 250 μM, and an invariant proteome, indicating a safety profile markedly superior to that of f1-C5C2.

**Figure 7.**
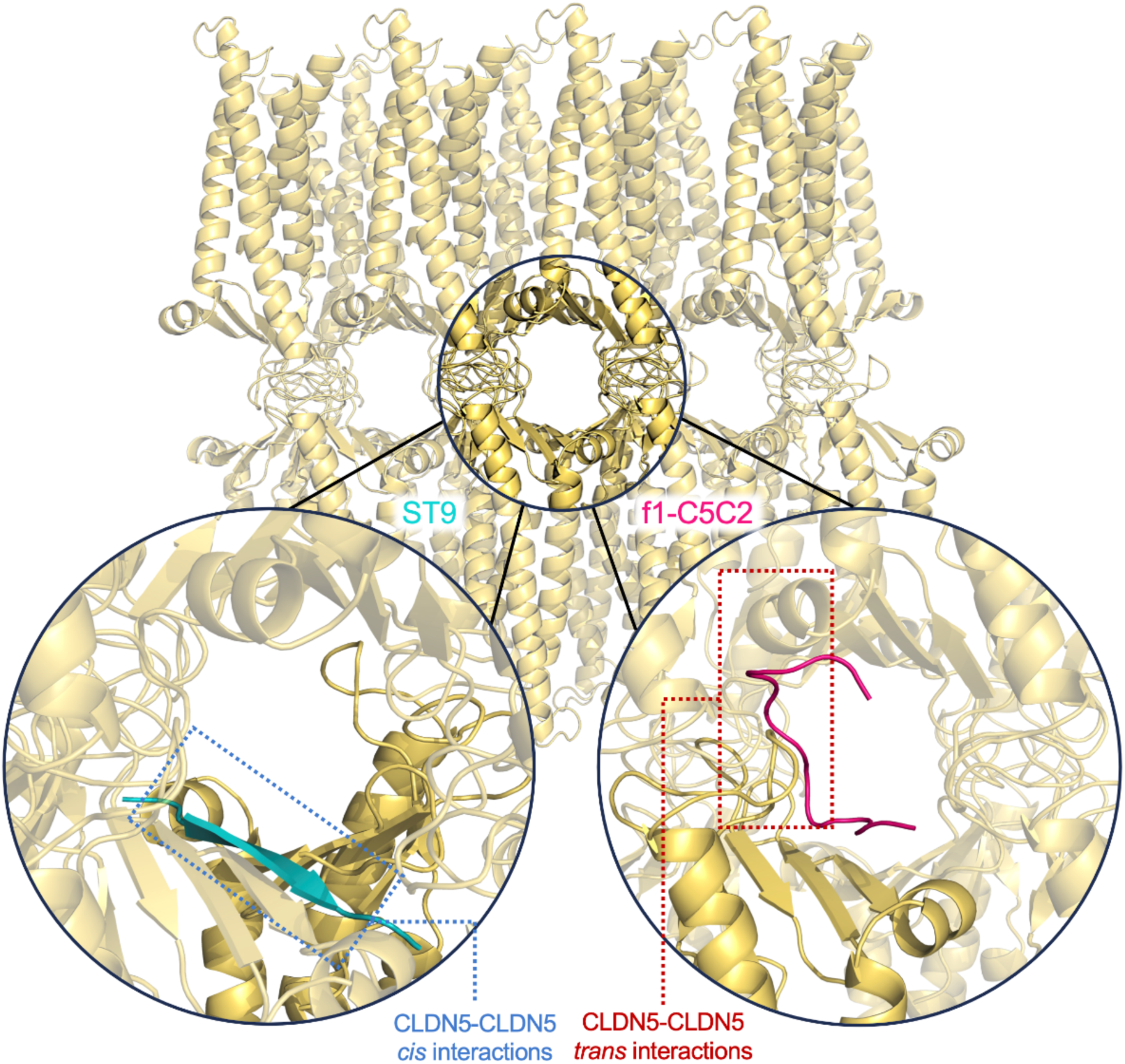
Putative binding sites for f1-C5C2 and ST9 in the multi-pore CLDN5 assembly. ST9 binds the protein at its *β*-sheet domain, potentially interfering with the CLDN5-CLDN5 *cis*-interactions at the basis of the *β*-barrel stabilization in the pore scaffold. In contrast, f1-C5C2 establishes contacts with the unfolded region of ECL2, inhibiting CLDN5-CLDN5 *trans*-interactions between strands from opposing cells. The multi-pore model is based on the structure reported in Ref.^11^.

In view of clinical translation, a particularly relevant application of controlled BBB opening is the restoration of adequate cerebral glucose supply in GLUT1DS. ST9 pretreatment significantly increases the paracellular translocation of D-glucose across bEnd.3 monolayers starting from 2 h of exposure. Critically, this increase occurs in the absence of any detectable upregulation of GLUT1, indicating that enhanced glucose transport is attributable exclusively to the opening of the paracellular pathway rather than to compensatory transcellular mechanisms, a distinction that could not be established for f1-C5C2, for which a contribution of GLUT1 upregulation could not be excluded.

In conclusion, the integration of ML generative algorithms into our computational workflow represents a significant step forward in the design of clinically relevant peptides targeting CLDN5 at TJ strands. ST9 achieves nanomolar affinity for CLDN5, induces a rapid, size-selective, and fully reversible increase in BBB paracellular permeability, preserves TJ protein expression and localization, and leaves the endothelial cell proteome unaltered, an unprecedented profile among reported CLDN5-targeting peptides. Future refinement of the design protocol through the adoption of conditioned generative models^33,38,60^, in which the diffusion process is guided by explicit selectivity constraints against structurally related off-targets such as CLDN1, CLDN2, and CLDN12, co-expressed at the BBB and sharing a conserved ECL fold with CLDN5, may further improve isoform specificity and reduce the risk of off-target effects. *In vivo* validation will constitute the critical next step, providing a more realistic assessment of ST9’s biological performance in living systems. This will help establish the translational basis for using next-generation BBB-penetrant agents to treat GLUT1DS and, more broadly, for delivering therapeutic molecules to the CNS across a range of neurological diseases.

## MATERIALS AND METHODS

### Protocol to design *de novo* claudin-5 peptide binders

To design, optimize, and rank candidates CLDN5 binders, we leveraged deep-learning-based architectures tailored for biologically relevant protein design and structure prediction.^33,38^ This pipeline was integrated with biocompatibility predictions (water solubility and toxicity) and MD simulations to assess complex stability and unbinding FE (**Figure 8**). First, we applied the generative diffusion model RFDiffusion^33^ to design peptide binder backbones against the CLDN5 target. Then, for each backbone candidate, the deep learning protein sequence design model ProteinMPN^34^ was used to predict the optimal sequence. Finally, the list of designed sequences was filtered based on structural optimization and ranked using AF^35^. The highest-ranked sequences obtained through this workflow were then further screened by predicting their biocompatibility through *in silico* assessment of their water solubility and toxicity. A detailed description of these steps is provided in the **Supplementary Computational Methods** Section.

**Figure 8.**
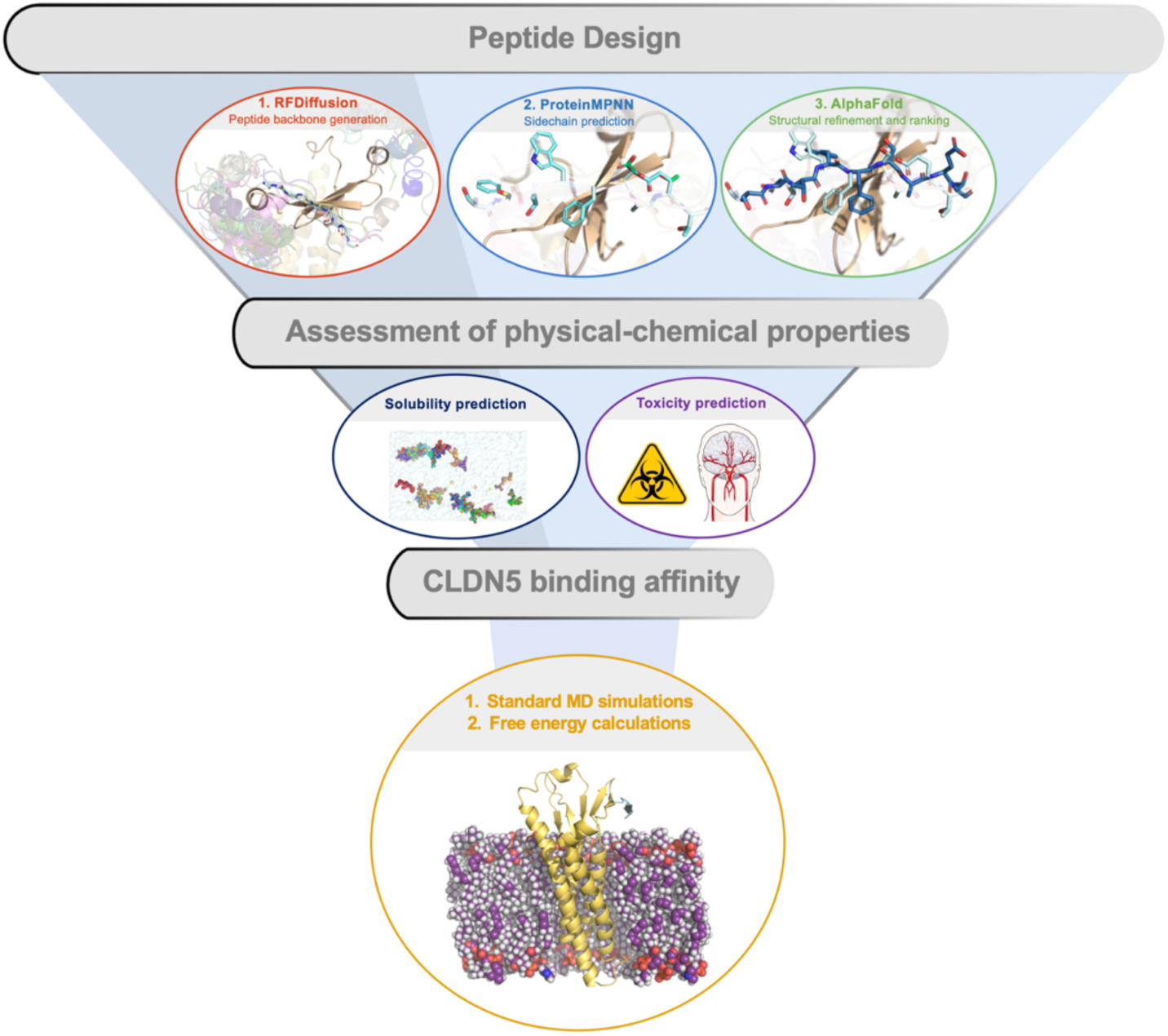
Schematic workflow of peptide design and computational characterization. Initial peptide design was carried out using a computational pipeline comprising RFdiffusion, ProteinMPNN-FR, and AF2, generating 4000 candidate sequences ranked according to their *pAE_interaction* scores. The three top-ranked peptides (GT9, ST9, and SA14) were subsequently evaluated *in silico* for solubility and cytotoxicity. ST9 and SA14 were selected for MD simulations to assess their binding to CLDN5. Finally, the binding affinity of ST9 was quantitatively estimated through FE calculations and compared with that previously reported for f1-C5C2 in Ref.16.

The two sequences that satisfied the screening criteria (ST9 and SA14) were tested for binding affinity to CLDN5 proteins via MD simulations and FE calculations, yielding a single best candidate, which was selected for experimental characterization. Details regarding the CLDN5-peptide binding affinity analysis are provided below.

### Computational assessment of claudin5-peptide binding affinity

#### Setup of the CLDN5-peptide complexes

The refined poses resulting from the AF2 optimization were used as the initial conformation for simulating the CLDN5-peptide complexes. Following the same protocol used in our previous work^16^, this system was embedded in a rectangular 1-palmitoyl-2-oleoyl-sn-glycero-3-phosphocholine (POPC) lipid bilayer with an XY area of 75 x 75 Å^2^, solvated with TIP3P^61^ water and a 0.15 M KCl ionic bath, using CHARMM-GUI^62^. The resulting systems included ∼ 68,000 atoms.

#### All-atom MD simulations

After a preliminary minimization and 30 ns of equilibration, 500 ns of MD simulations were performed with NAMD 3.0^63^ and the CHARMM36m force field^64^ in the NPT ensemble at T=310 K and P=1 bar maintained by a Langevin thermostat and the Nosé-Hoover Langevin piston pressure control^65,66^ (CHARMM-GUI default parameters were used). During production, the protein was restrained from lateral movement by imposing weak positional restraints to the C*α* atoms of the TM residues Leu13, Ala82, Val93, Thr120, Leu126, Cys137, Trp168, Val176. The disulfide bond between residues Cys54 and Cys64 found in the ECL1 domain was preserved. Rectangular Periodic Boundary Conditions (PBCs) were used to replicate the system and remove box surface effects. Long-range electrostatic interactions were computed using Particle Mesh Ewald (PME)^67^. Electrostatic and VdW interactions were calculated with a cutoff of 12 Å as prescribed by the CHARMM force field. A switching function was applied, starting to take effect at 10 Å to obtain a smooth decay^68^. Hydrogen atoms not involved in covalent bonds with water were restrained with SHAKE^69^, while those of water molecules were kept fixed with SETTLE^70^, enabling the use of a timestep of 2 fs.

Three independent replicas were produced for each system, and the results and associated error of each analysis are expressed as the mean and standard deviation over the three replicas.

#### Free energy calculations of CLDN5–peptide unbinding

Due to the stability of the CLDN5– peptide complexes, peptide dissociation events are rarely observed in conventional MD simulations. Therefore, enhanced sampling techniques were employed to characterize the unbinding process and to obtain reliable estimates of the relative FE. Consistent with Ref.16, enhanced sampling was performed with temperature-accelerated Molecular Dynamics (TAMD) simulations^40,41^. Briefly, during TAMD, a set of CVs, functions of the system’s atomic coordinates, is coupled to auxiliary fictitious variables that evolve at an artificial temperature higher than the physical system’s temperature. This temperature difference allows the slow degrees of freedom (CVs) to diffuse across FE barriers, effectively dragging the underlying atomic coordinates with them. To avoid heat transfer to the physical system, a separation of timescales is introduced, so that the auxiliary CVs evolve more slowly than the physical atomic coordinates. The method enables the reconstruction of FE landscapes by integrating the forces acting on the CVs, which are, by definition, the derivatives of the FE with respect to the CVs, also called the mean forces^16,40,42^. For additional details on the method and the underlying theory, we refer the reader to Refs ^40,41^.

#### TAMD implementation

We selected, as a CV, the distance between the centers of mass (COMs) of the peptides and the CLDN5 ECL, mapped in a range from 8 to 40 Å. To achieve faster exploration of the CV-space, this was split into three independent windows, from 8 to 17 Å, from 17 to 30 Å, and from 30 to 40 Å. Each window was simulated for 500 ns. An effective temperature 1000 K and an effective friction coefficient of 15 ps^-1^ were used for the CVs dynamics, and the instantaneous mean forces were collected in bins of 0.1 Å. To ensure a faster convergence of the calculations, the lateral diffusion of the peptide was limited by a cylindrical restraint^71–76^.

TAMD was implemented using the Colvars module ^77^, via the extended Lagrangian dynamics feature for the CVs^42^, and the FE was computed from the mean forces using the corrected z-averaged (CZAR) estimator^78^. The TAMD runs were carried out with the same MD set-up of the standard simulations previously described. The FE profile and the associated error were reported as the mean and the standard deviation from three independent replicas, respectively.

### Cell culture and transfection procedures

Mouse brain endothelial bEnd.3 cells (ATCC® CRL-2299™) were obtained from ATCC and expanded in DMEM containing 10% fetal bovine serum (FBS), 1% penicillin-streptomycin (PS), and 1% glutamine (G). Cultures were kept at 37 °C in a humidified incubator with 5% CO₂, and the culture medium was refreshed every two days. Cells were used between passages 25 and 30. For experiments requiring endothelial monolayers, bEnd.3 cells were plated either on collagen-coated Transwell® inserts (150 µg/mL collagen; 12 mm diameter; 0.4 µm pore size; 1.12 cm2 growth area; Transwell 3460, Corning® Costar®) at a density of 40 × 103 cells/insert, or on the appropriate culture support, including Petri dishes, glass coverslips, or black 96-well plates, depending on the downstream assay. Unless otherwise indicated, reagents were purchased from ThermoFisher Scientific.

HEK293T cells (ICLC, HTL04001), which constitutively express the SV40 large T antigen, were maintained under the same culture conditions, using DMEM supplemented with 10% FBS, 1% PS, and 1% G. For transient expression of mouse CLDN5, cells were seeded on poly-D-lysine-coated Petri dishes (10 μg/mL) 24 h before transfection. At approximately 80% confluence, cells were transfected with 40 μg of plasmid DNA encoding mouse CLDN5 fused to tGFP (mCLDN5-tGFP, Origene #MG202442). Lipofectamine 2000-based transfection complexes were prepared in DMEM, incubated at room temperature (RT) for 20 min, and then added to the cells. After 6 h of incubation at 37 °C, the transfection medium was replaced with fresh complete medium. Cells were collected 24 h later for lysis and subsequent binding assays. Fluorescent protein content in the resulting lysates was estimated from absorbance at 280 nm using a NanoDrop 2000 spectrophotometer. The amount of tGFP present in the lysates was determined by comparison with a calibration curve generated using recombinant tGFP from *Pontellina plumata* (Origene #TP700079). Fluorescence measurements were performed with a Tecan SPARK multimode microplate reader (Tecan, Switzerland).

### Microscale thermophoresis binding assay

Lysates from HEK293T cells expressing mCLDN5-GFP were generated according to our previously reported protocol^16^ and used as fluorescent target material for microscale thermophoresis (MST) experiments. Peptides (ST9 and SA14) were prepared in PBS/0.1% Tween-20 as 15-point two-fold serial dilutions, with a maximum concentration of 200 μM, and incubated with lysates containing 1 μM mCLDN5-GFP. Samples were loaded into Premium capillaries (Monolith Series, #MO-K025, NanoTemper) and analyzed using a Monolith NT.115 instrument with the Nano-BLUE laser set to 20–60% power. Binding curves were fitted to determine the apparent dissociation constant Kd. Primary analysis was performed with MO.Affinity Analysis software using a 1:1 interaction model. Because peptide binding induced changes in initial fluorescence, the effect on normalized fluorescence (Fnorm) was corrected using DI.Screening Analysis software. K_d_ values were additionally verified by fitting the binding curves in GraphPad Prism 10 using a one-site-specific binding nonlinear regression model, which yielded comparable estimates.

### TEER measurements and transport studies

Cells were cultured on Transwell inserts until TEER values reached a stable plateau of ∼10-15 Ω·cm². The apical medium was then replaced with medium containing the indicated treatment or vehicle. For ST9 experiments, cells were treated with ST9 (100-250 μM) or vehicle. To assess the contribution of autophagy, cells were treated with f1-C5C2 (25 μM) in the absence or presence of rapamycin (100 nM). In addition, the effect of nutrient deprivation was evaluated by culturing cells in medium supplemented with 2% FBS for 24 h before incubation with peptide or vehicle. TEER was recorded hourly for 72 h using a CellZscope+ system (NanoAnalytics). Paracellular transport was quantified by measuring the basolateral accumulation of FITC-dextran (4, 40, or 70 kDa; FD4, FD40, FD70; Merck) at selected time points. Transport studies were performed using supplemented FluoroBrite™ DMEM medium (A1896701, Gibco, Thermo Fisher Scientific). Before each measurement, the apical compartment was replaced with FluoroBrite™ DMEM containing FITC-dextran (40 μg mL-1) and the corresponding treatment, while the basolateral compartment was filled with 1.5 mL of fresh FluoroBrite™ DMEM. At each time point, 100 μL was collected from the basolateral chamber into a 96-well plate and immediately replaced with 100 μL fresh medium to maintain constant volume. Fluorescence was measured using a Tecan SPARK multimode microplate reader (excitation 485 nm; emission 535 nm). Calibration curves were generated from serial dilutions of FITC-dextran (0-40 μg/mL) prepared in FluoroBrite™ DMEM, and linear regression was used to convert fluorescence values into dextran concentration and to calculate the total basolateral mass. Fits typically yielded r^2^ = 0.98-0.99 (n = 3 wells from three independent culture preparations). The translocation rate was calculated as the ratio of the basolateral dextran mass at each time point to the initial apical mass (t = 0) and normalized to transport across cell-free Transwell inserts, expressed as a percentage of the no-cell control.

### D-Glucose transport studies

Transport studies were performed by measuring D-Glucose (G8270, Sigma-Aldrich) permeability in bEND.3 brain endothelial cells at 37 °C at 2 and 8 h. Once reaching confluence, cells on Transwell® were treated with either 25 µM f1-C5C2 or 250 µM ST9 for 24 h. In particular, the apical fraction was filled with complemented FluoroBrite™ DMEM cell medium, while non-complemented glucose-free DMEM (A1443001, Gibco, ThermoFisher Scientific) was used for the basolateral compartment to avoid interference with the following experimental step. The day after, cells were washed once with PBS prior to incubation with FluoroBrite™ DMEM containing 10 mg/mL, and the basolateral compartment was filled with 1.5 mL of fresh glucose-free medium. At the indicated time points, 50 µL aliquots were sampled from the basolateral chamber and immediately replaced by 50 µL of fresh cell medium to maintain the total volume. Glucose quantification was performed using the Glucose Colorimetric Detection Kit (EIAGLUC, Invitrogen, ThermoFisher Scientific). Basolateral fractions were diluted 1:30 in the kit Assay Buffer prior to use. Results were expressed as normalized translocation rate, as in dextran transport studies.

### Cell viability and caspase 3/7 activation

To assess apoptotic and necrotic cell death, bEnd.3 cells were treated with selected peptides (25 μM f1-C5C2 or 250 μM ST9) for 4 h. To investigate the contribution of autophagy, cells were exposed to f1-C5C2 (25 μM) in the absence or presence of rapamycin (100 nM). The effect of nutrient deprivation was evaluated by culturing cells in medium supplemented with 2% FBS for 24 h before peptide or vehicle treatment. After one wash with PBS, cells were incubated for 30 min with CellEvent™ Caspase-3/7 Green Detection Reagent (Invitrogen, C10423) to detect apoptotic cells. Cells were then counterstained for an additional 5 min with Hoechst 33342 (1 μM) for nuclear visualization and propidium iodide (PI, 1 μM) for necrotic cell detection. Images were acquired using a Nikon Eclipse 80i upright epifluorescence microscope at 10× magnification. Cell death quantification was performed on 10 fields per sample from three independent cultures using CellProfiler software (Broad Institute).

### Western blotting of TJ proteins

bEND.3 cells were seeded in six-well plates and grown to confluence, then treated with the ST9 peptide at 250 μM concentration (or 25 μM f1-C5C2). At 24 h, cells were washed three times with ice-cold PBS and lysed in RIPA buffer (50 mM Tris-HCl pH 7.4, 150 mM NaCl, 1% Igepal, 0.1% SDS, 0.5% sodium deoxycholate) supplemented with EDTA-free protease inhibitors (Roche Diagnostic) and serine/threonine and tyrosine phosphatase inhibitor cocktails (Sigma-Aldrich). Lysates were collected by scraping, briefly sonicated (10 s; Branson SLPe, 25% amplitude) and centrifuged (19,000g, 15 min, 4 °C). Protein concentration was determined by BCA assay (Thermo Fisher Scientific) using a BSA standard curve. Samples were denatured (98 °C, 5 min), resolved by SDS-PAGE (5% stacking gel; 8-12% resolving gels) and transferred to nitrocellulose membranes (Amersham Protran, Cytiva) (100 V, 90 min, 4 °C). Membranes were blocked in 5% milk in TBS-T (0.05% Tween-20) for 1 h at room temperature and incubated overnight with primary antibodies against ZO-1 (Invitrogen, #61-7300, 1:500), Occludin (Invitrogen, #33-1500, 1:1000), claudin-5 (Invitrogen, #PA599415, 1:500), GLUT1 (Novusbio, NB110-39113, 1:1000), and calnexin (Enzo Life Sciences, ADI-SPA-860, 1:1000). After three washes in TBS-T, membranes were incubated with HRP-conjugated secondary antibodies (Abcam, 1:10000) for 1 h at room temperature and developed using ECL Prime Western Blotting System (Cytiva). Chemiluminescence was acquired on an iBright FL1500 imaging system (Thermo Fisher Scientific, A44241), and band intensities were quantified with iBright Analysis Software.

### Immunofluorescence

bEnd.3 cell monolayers were grown on black 96-well plates (μ-Plate 96 Well Square, Ibidi, #89626) and exposed to the indicated treatments. Cells were fixed with 4% paraformaldehyde (PFA) in PBS for 15 min at room temperature (RT), permeabilized with 0.1% Triton X-100 for 5 min, and blocked with 2% BSA in PBS for 30 min. Samples were then incubated with anti-CLDN5 primary antibody (Invitrogen, #PA599415, 1:500) diluted in blocking buffer at RT for 3 h. After repeated PBS washes, cells were incubated for 1 h at RT with species-appropriate secondary antibodies (Thermo Fisher Scientific, 1:500) diluted in blocking buffer. Nuclei were counterstained with Hoechst 33342 (1 μM; Sigma-Aldrich, #3342) for 5 min, and samples were mounted with Vectashield antifade medium (Vector Laboratories, #H-1000-10). Confocal images were acquired using a Leica SP8 laser-scanning microscope with a 63× objective. Within each experiment, acquisition settings were kept constant across all conditions. For each sample, five fields of view were acquired as matched z-stacks. Maximum-intensity projections were generated and analyzed in CellProfiler using a custom pipeline, as previously described^16^.

### Cell proteomics and bioinformatic analysis

#### Cell lysate preparation for proteomic analysis

For proteomic analysis, bEnd.3 cells were cultured in six-well plates and exposed at confluence to ST9 (250 μM) or f1-C5C2 (25 μM) for 24 h, following the same treatment and lysis procedure described for western blotting. Briefly, cells were lysed in RIPA buffer supplemented with protease and phosphatase inhibitors, and clarified lysates were quantified by BCA assay before downstream proteomic processing.

#### Protein digestion and MS/MS analysis

50 ug of proteins in RIPA buffer were processed as follows: cysteine residues were reduced and alkylated with a final concentration of 10 mM TCEP and 30mM Chloroacetamide (95 °C for 15 min in the dark). After cooling to RT, samples were digested according to the SP3 procedure^79^. The concentration of the resulting peptides was determined using a Fluorometric Peptide Assay (ThermoFisher), and 300 ng of peptides were used for each proteomic analysis. Proteomic analysis was conducted on a Thermo Exploris 480 orbitrap system coupled with a Dionex Ultimate 3000 nano-LC system. After trapping and desalting, the tryptic peptides were loaded on an Aurora C18 column (75 mm x 250 mm, 1.6 µm particle size) nanocolumn (Ion Opticks, Fitzroy, Australia) and separated using a linear gradient of acetonitrile in water (both added with 0.1% formic acid), from 3% to 41% in 1 h, followed by column cleaning and reconditioning. The flow rate was set to 300 nL/min. Total run time was 1.5 h. Injection volume was set to 1 µL. Peptides were analyzed in positive ESI mode with a capillary voltage of 2.0 kV. The RF lens was set to 40%, and the AGC target set at 300%. Data acquisition was performed in Data-Independent mode (DIA) with a survey scan set from 400 to 1000 m/z at 120,000 resolution, followed by MS/MS acquisition of 60 m/z transmission windows, each with a fixed 10 Da width. MS/MS spectra were acquired in HCD mode at 15,000 resolution. All the collected MS/MS spectra were analyzed using Spectronaut (Version 18), by running a DirectDIA analysis against the reference *Mus musculus* FASTA database (Tax ID: 10090 reporting 17228 reviewed entries). Cysteine carbamidomethylation was selected as a fixed modification; acetylation of protein N-Term and Methionine oxidation were selected as variable modifications. Positive protein identifications were retained at 1% false discovery rate (FDR) threshold, and at least two peptides were used for protein quantification.

### Differential Expression Analysis

Data analysis was performed in the R environment, version 4.3.3. For each sample, three technical replicates were aggregated by calculating their average abundance. Subsequently, only proteins whose values were quantified across all analyzed samples were retained.

The DESeq2 algorithm^80^ was utilized to evaluate differential protein expression between the ST9 treated samples and controls and between the f1-C5C2 treated samples and controls. To reduce noise associated with low or highly variable count estimates, LFC shrinkage was applied. Proteins were considered significantly differentially expressed if they exhibited an absolute fold change greater than 1.5 and an adjusted p-value smaller than 0.05.

### Functional Annotation and Gene Set Enrichment Analysis (GSEA)

To characterize the functional significance of the variations in protein abundance, Gene Set Enrichment Analysis (GSEA) was performed using the clusterProfiler R package^81^. Enrichment was evaluated for the Biological Process (BP), Molecular Function (MF) and Cellular Component (CC) Gene Ontology categories using the gseGO function of clusterProfiler. The p-values computed for each pathway were corrected using the Benjamini & Hochberg method, and only adjusted p-values < 0.05 were considered significant.

## Statistical analysis

All statistical analyses were performed with GraphPad Prism v10. Results are reported as mean ± SEM, and individual data points are shown in the graphs when appropriate. The number of independent biological replicates is specified in each figure legend. Data distribution was evaluated using the D’Agostino-Pearson normality test, when applicable. For comparisons between two groups with normal distribution, a two-tailed Student’s *t*-test was applied. When more than two normally distributed groups were compared, one-way or two-way ANOVA was used according to the experimental design, followed by Tukey’s post hoc test for multiple comparisons. For time-course or matched datasets, repeated-measures ANOVA was applied. Non-normally distributed matched datasets were analyzed using Friedman’s test followed by Dunn’s multiple-comparisons test. Statistical significance was set at p < 0.05.

## Acknowledgements

We are grateful to the Fondazione Istituto Italiano di Tecnologia High-Performance Computing Infrastructure (Genova, Italy), S. Decherchi for useful discussions about machine learning, Diego Moruzzo and Arta Mehilli for technical help, and Dr. Rossana Ciancio for administrative assistance (Istituto Italiano di Tecnologia, Genova, Italy). We acknowledge ISCRA for awarding this project access to the LEONARDO supercomputer, owned by the EuroHPC Joint Undertaking, hosted by CINECA (Bologna, Italy).

## Author Contributions

Authors’ contributions are attributed on the basis of CRediT author statement. Conceptualization: VC, LM, FB. Methodology: AB, MT, GA, AP, ADF. Software: AB, GA, LM, AP. Validation: AB, MT, AA, AP, VC, LM, FB. Formal analysis: AB, MT, GA, AP, ADF, AA, VC, LM, FB. Investigation: AB, MT, GA, AP, ADF, AA, VC, LM, FB. Resources: LM, FB. Data curation: AB, MT, AA, VC, LM, FB. Writing - original draft: AB, VC, LM, FB. Writing - review & editing: AB, MT, GA, AP, ADF, AA, VC, LM, FB. Visualization: AB, MT, AP, VC, LM, FB. Supervision: VC, LM, AA. Project administration: VC, LM, FB. Funding acquisition: VC, LM, FB.

## Data, code, and materials availability

All data and code needed to evaluate and reproduce the conclusions in the paper are present in the paper and/or the Supplementary Materials. Structural models and MD simulations data (input files, parameters, and starting configurations) are available at the following doi: 10.5281/zenodo.21456066. The NAMD code is publicly available at: https://www.ks.uiuc.edu/Research/namd/.

