## Supplementary material for "Controlled Blood-Brain Barrier Modulation by a High-Affinity Claudin-5 Peptide Binder": SI

**Short title:**
Peptide modulation of blood-brain barrier permeability

[‡] These authors have contributed equally.
[§] Corresponding authors
^ Present address: Alessandro Berselli, Atomistic Simulations, Center for Human Technologies, Istituto Italiano di Tecnologia, 16156 Genova, Italy. Martina Trevisani, Translational Cancer Medicine Program, 00014 University of Helsinki, Finland.

### SUPPLEMENTARY INFORMATION

### Supplementary Computational Methods

#### Peptide design

1) *Backbone generation with RFDiffusion.*

RFDiffusion is a generative diffusion model that generate highly realistic and reasonable protein structures with specific geometries through iterative denoising trajectories originating from an initial random noise sample. This approach proved effective in generating protein monomers, multimers, functional protein and enzymatic scaffolds and protein binders[33,36,55,56,60]. In particular, we selected as a protein target the configuration of the CLDN5 monomer ECL domain extracted from the equilibrated multi-pore model[11], comprising residues 28 to 77 (ECL1) and 149 to 161 (ECL2). To drive structure generation towards attractive potential binding sites, the model was minimally conditioned by specifying two hydrophobic residues (Val70 and Val154) as *hotspots*. During denoising, these residues are encouraged, but not forced, to be contacted by predicted structures[33], and these were selected as they were the most conserved binding sites observed in our previous work[16].

We designed 1000 structures with length comprised between 5 and 15 residues, applying denoising scaling factors of 0.5 to both C$\alpha$ coordinate and backbone geometry.

2) *Side chain prediction with ProteinMPNN*

For each structure predicted with RFDiffusion, the most likely sequences correlating the given folding were predicted with the message-passing neural network model ProteinMPNN[34]. This model uses as an input the backbone structure of the protein target, that is kept fixed trying to maximize the probability that a certain sequence produces the given folded geometry. This approach demonstrated an experimental success design almost 5-fold greater than Rosetta-design[38]. However, it assumes that the peptide backbone remains static, thus leading to steric clashes or sub-optimal structural prediction. For this reason, sequence prediction by ProteinMPNN is combined by structural refinement with the FastRelax (FR) algorithm, enabling minimal structural optimization through side chain repacking and backbone adjusting (https://github.com/nrbennet/dl_binder_design).

For each of the 1000 backbones obtained from RFDiffusion we generated four sequences, resulting in a total of 4000 structures.

3) *Structural optimization and ranking with AlphaFold.*

Following the protocol described in Ref.s 38,82, the experimental success design was evaluated based on the metrics provided by AF2 scoring. In particular, the predicted aligned error (pAE) is the AF2 outputs evaluating the confidence of the relative position for each pair of residues in the predicted model, resulting in a $N \times N$ matrix, with $N$ being the number of residues. By summing the values only for the inter-chain blocks of the pAE matrix, the *pAE_interaction* score is obtained, that estimates the confidence about the relative orientation and placement of two interaction chains. According to the benchmarks reported in Ref.s 38,82, designs with AF2 *pAE_interaction* < 10 Å were assigned an experimental low nanomolar binding affinity ($K_d$).

The 4000 sequences were refined and filtered with AF2 and ranked according to the resulting *pAE_interaction* score. After careful analysis of the results, a cutoff of 13 Å was chosen to promote sequences for subsequent analysis.

#### Computational screening based on physical-chemical properties

From the AF2 filtering, three sequences were selected for deeper analysis, which were named GP9, SA14 and ST9. Before proceeding with detailed investigation of the CLDN5-binding affinity, we tested their solubility and toxicity in the water-based cell medium, as described below.

1) *Solubility prediction*

To provide a precise prediction of the solubility in the cell medium, we based our calculation on the computational protocol optimized in our previous work[16] and inspired by Ref.83

**Simulation set-up.** The procedure consists of assembling a grid of 3 x 3 x 3 equally spaced, non-interacting copies of the peptide of interest, solvated in a water simulation box and a mixed KCl/$CaCl_2$ ionic bath mimicking the same ionic strength as that used for *in vitro* experiments. The N- and C-termini of each peptide were neutralized by end-capping with amide and ester groups, respectively. After short equilibration with peptide backbones fixed, this system underwent 100 ns of standard MD simulations, capturing the tendencies of the peptides to aggregate or remain hydrated.

**Analysis.** To quantify the extent of peptide aggregation during MD simulations we calculated three descriptors: (i) the size of the largest aggregate (in number of peptides), the number of different aggregates and the average water-peptide contact number expressed as a percentage with respect to the initial configuration. These descriptors were calculated over the last 10 ns of trajectories. Full details are reported in Ref.16.

**Solubility Classification.** The classification of peptides into either *soluble* or *insoluble* categories based on the reported descriptors would be ambiguous in the absence of a reference. For this reason, we replicated the same protocol using an external dataset of 16 sequences selected from the Antimicrobial Peptide Database (https://aps.unmc.edu ) based on their hydrophilicity. Specifically, we identified 8 hydrophilic (hydrophilic residues > 70%) and 8 hydrophobic (hydrophilic residues < 20 %) peptides, which were used to train a supervised learning classification model based on the K-Nearest Neighbor (KNN) algorithm (k=3, uniform weight function). The average dimension of the largest aggregate, the number of aggregates, and the percentage of water contacts were used as independent variables to train the model.

2) *Toxicity prediction*

The toxicity of the potential CLDN5-binders was predicted with ToxinPred (https://webs.iiitd.edu.in/raghava/toxinpred3)[39], using the default scoring threshold 0.38.

### Supplementary Results

#### AlphaFold scoring of designed peptides.

As described in the Computational Methods, we generated a library of 4,000 peptide sequences using RFDiffusion followed by ProteinMPNN-FR design and subsequently refined the peptide–CLDN5 binding interfaces using AF2.

In contrast, the complex RMSD, which quantifies the average atomic deviation between the predicted structure and the design model, and the predicted Local Distance Difference Test (pLDDT), which provides a per-residue confidence score for the structural model report primarily the accuracy with which the peptide fold as designed. Since our primary objective is to identify peptides capable of binding CLDN5, we ranked the designed sequences by their *pAE_interaction* scores.

The distributions of the RMSD of the peptides refined with AF2 with respect to the structures designed with RFDiffusion (**Figure S2A**) shows that most structures exhibit values < 1.5 Å, indicative of low divergence between the two models. In contrast, greater variability is observed in the pLDDT scores (**Figure S2B**), which estimate the confidence in AF2's structural prediction. The distribution spans from 40 to 90, with two peaks at ~55 and ~75, suggesting the presence of two main populations of structures: a predominant one characterized by low confidence and likely structural flexibility and unfolded domains, and a second with substantially higher confidence, reflecting a well-defined fold. High values of pLDDT correlate with low values of RMSD, meaning that high structural confidence by AF2 is also predictive of agreement with RFDiffusion modeling.

In line with this evidence, the analysis of the secondary structure elements performed with pyPept[85] (**Figure S2D**), reveals a remarkable preference for unfolded structures, which represent more than 90% of the total designed peptides. The remaining sequences form mainly $\alpha$helical segments, which account for ~10% of the total structural domains, while the $\beta$sheet domains remain marginal, as they are predicted only for a few sequences.

However, the presence of defined structural elements does not imply higher confidence in the AF2 prediction. **Figure S2E** shows the binder RMSD versus binder pLDDT, colored by the percentage of helicity of each predicted sequence. The region associated with higher confidence (low binder RMSD and high binder pLDDT) does not preferentially correspond to sequences with greater helical content, indicating that structural definition alone is not a reliable indicator of predictive accuracy.

### Supplementary Figures

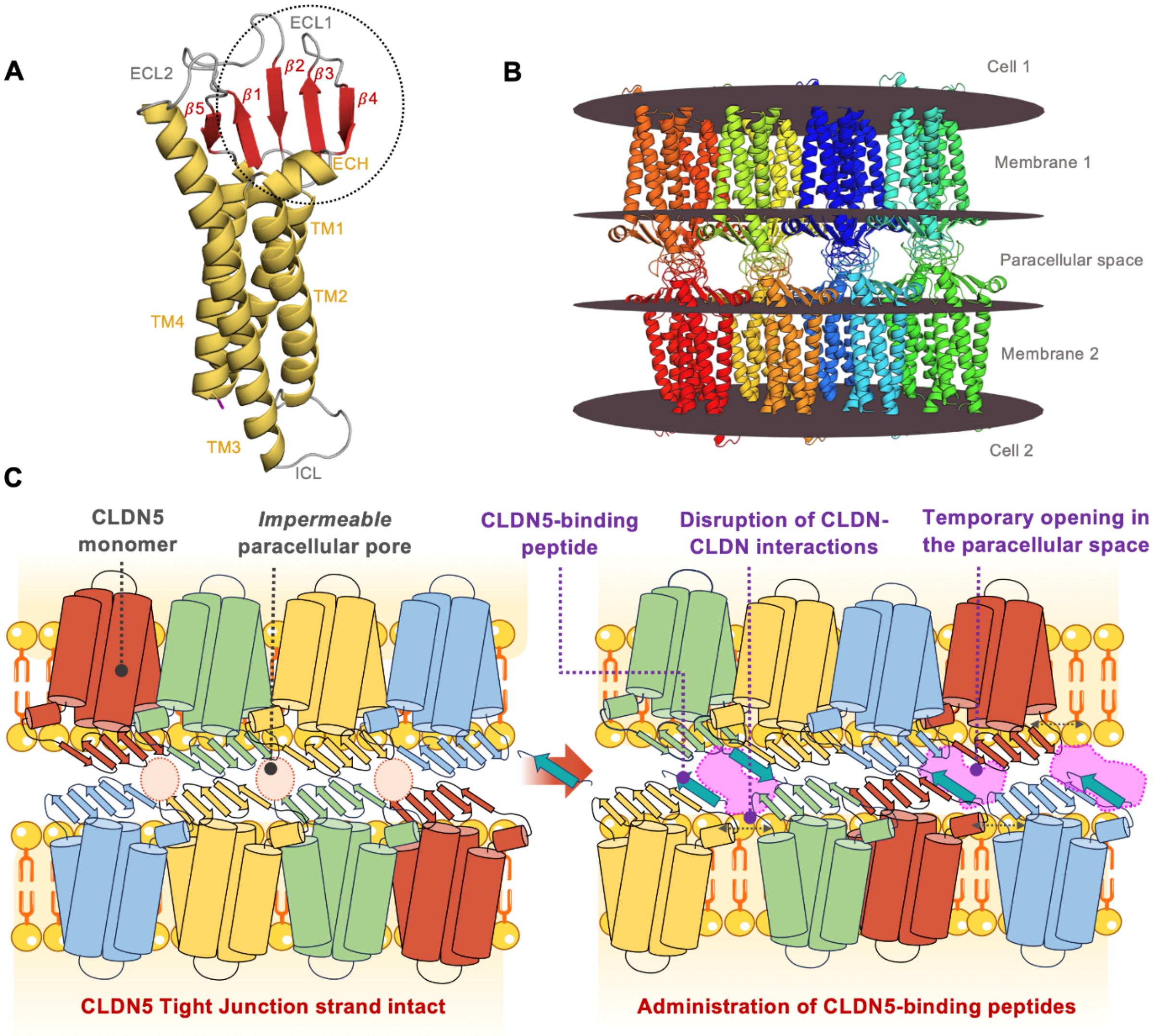

**Figure S1. Claudin-5 proteins in the Tight Junction strands.** (**A**) Three-dimensional structure of the CLDN5 monomer obtained via homology modeling starting from the CLDN15 crystal structure (PDB ID: 4P79)[86]. The protein is shown in cartoon representation, colored by secondary structure elements (yellow for $\alpha$-helices, red for $\beta$-sheets, and grey for unfolded domains). (**B**) Structural model of a representative three-pore segment of CLDN5 assembly within TJ strands (taken from Ref. 11). (**C**) Schematic representation of CLDN5 multimers in the intact TJ strand (*left*) and after administration of a CLDN5-targeting peptide (*right*). Upon binding to the ECL, the peptide disrupts CLDN5-CLDN5 interactions, creating transient openings across the paracellular space.

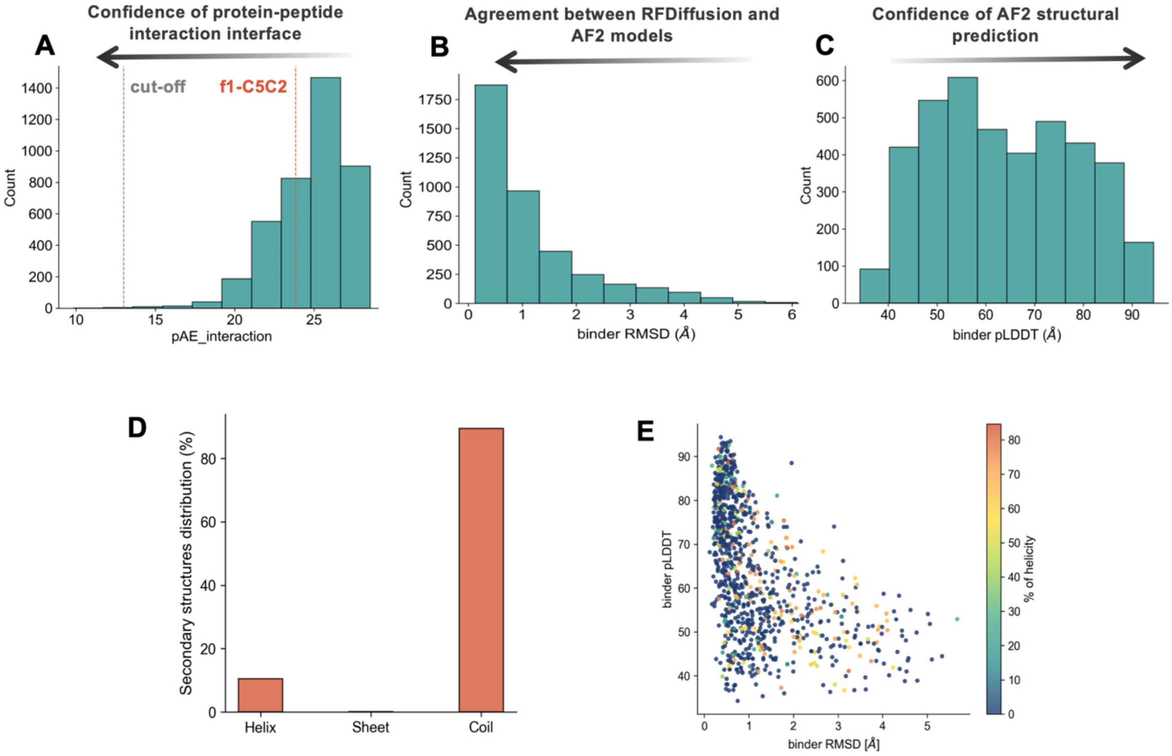

**Figure S2. Distributions of the scoring metrics and structural elements for the designed sequences as predicted by AlphaFold.** (**A**) The pAE_interaction reflects the confidence in the modeled protein–peptide interface. A cutoff value of 13 Å was applied to select sequences for subsequent analyses. (**B**) The binder RMSD was computed between the structures generated by RFDiffusion/ProteinMPNN and those predicted by AF2, providing a measure of structural consistency between design and prediction. (**C**) The binder pLDDT reports the confidence of the overall AF2 structural model. Arrows indicate the direction corresponding to increasing prediction accuracy for each metric. (**D**) The percentage of $\alpha$-helical, $\beta$-sheet, and random coil content across the ~4000 sequences was calculated using pyPept[85]. (**E**) Distribution of the AF2 metrics, binder RMSD versus binder pLDDT, for the designed peptides. Each point represents an individual peptide and is colored according to its percentage of helical content.

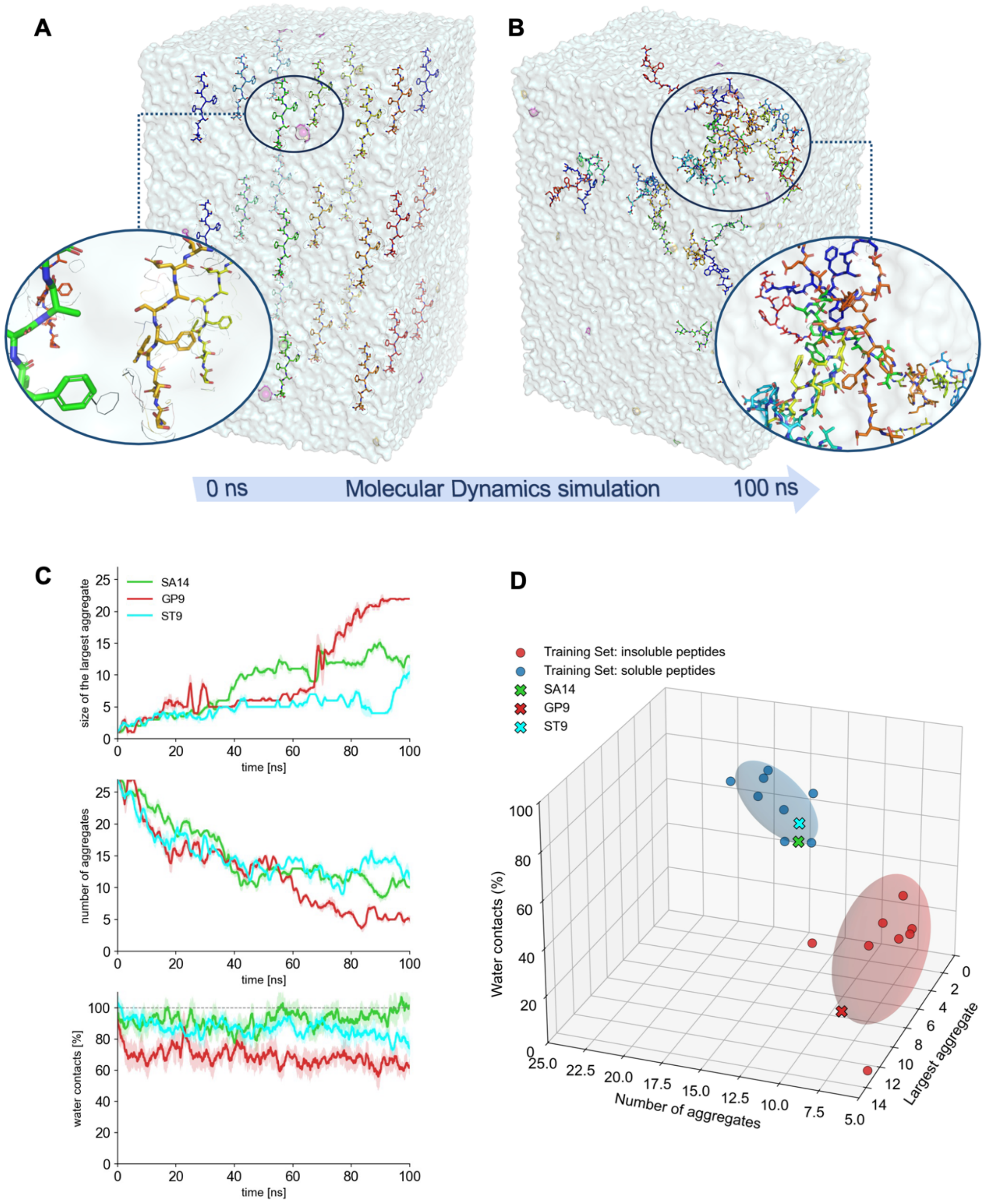

**Figure S3. Computational prediction of water solubility.** (**A**) Peptide solubility was evaluated by constructing a 3×3×3 grid of 27 equally spaced identical oligomers, capped with neutral N- and C-termini and solvated in water containing a mixed KCl/$CaCl_2$ ionic bath. For each peptide, an identical initial setup was generated, and the systems were simulated for 100 ns using standard molecular dynamics. (**B**) The aggregation state at the end of the simulation was quantified by calculating the number of aggregates, the size of the largest aggregate, and the percentage of total water contacts relative to the initial configuration. (**C**) Time trace of the descriptors for SA14 (green profile), GP9 (red profile), and ST9 (cyan profile). For clarity, traces report the moving average of the raw time series (window = 10 frames). Shaded bands indicate the local standard deviation computed within the same sliding window. (**D**) These descriptors were used to train a supervised k-nearest neighbors (KNN) model based on an external dataset of 8 soluble (blue) and 8 insoluble (red) peptides. The covariance ellipsoid is shown as a transparent surface. The three selected candidates (SA14, GP9, and ST9) are indicated by X symbols.

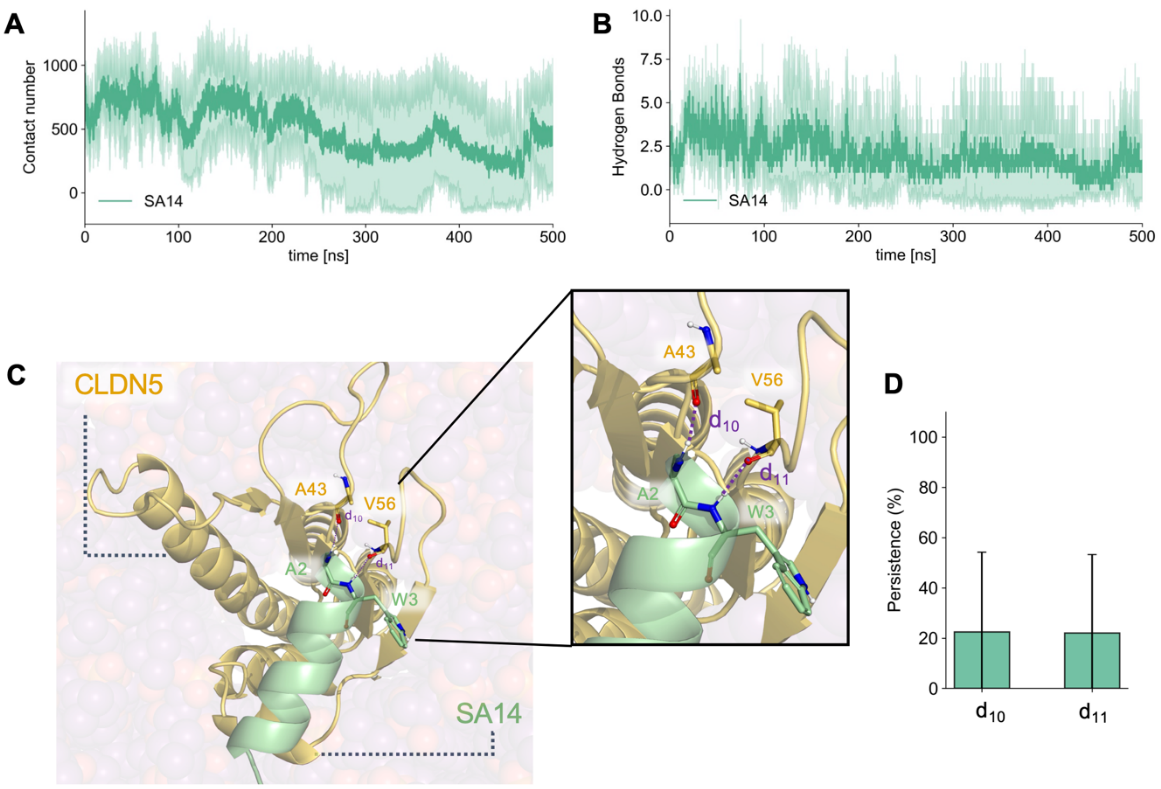

**Figure S4. Stability of the CLDN5-SA14 complex via molecular dynamics simulations.** Time evolution of the (**A**) contact number and (**B**) HBs between CLDN5 and SA14 during 500-ns-long MD simulations. (**C**) Interaction interface between CLDN5 ECL domain (yellow) and SA14 (green). The most persistent HBs detected during MD simulations are indicated with purple dotted lines. The protein and the peptide are shown with the cartoon representation. The oxygen, nitrogen, and hydrogen atoms involved in the interaction interface are reported as spheres and colored in red, blue, and white, respectively. (**D**) The persistence of HBs between CLDN5 and SA14 during MD simulations was calculated as a percentage of simulation time using a cutoff of 3.3 Å. Only interactions with persistence time > 20% were considered. The results and associated errors for each analysis are reported as the mean and standard deviation over three independent replicas.

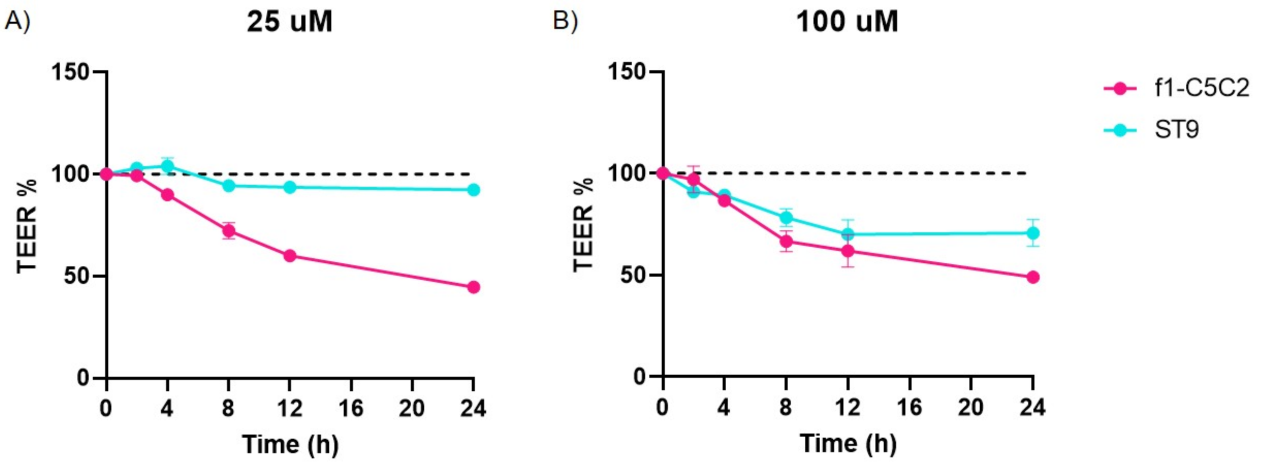

**Figure S5. Optimization of working concentration.** TEER values for bEND.3 cell layer (expressed in percent of the initial value) upon exposure to 25 µM (**A**) and 100 µM (**B**) of either f1-C5C2 (pink) or ST9 (cyan) peptides.

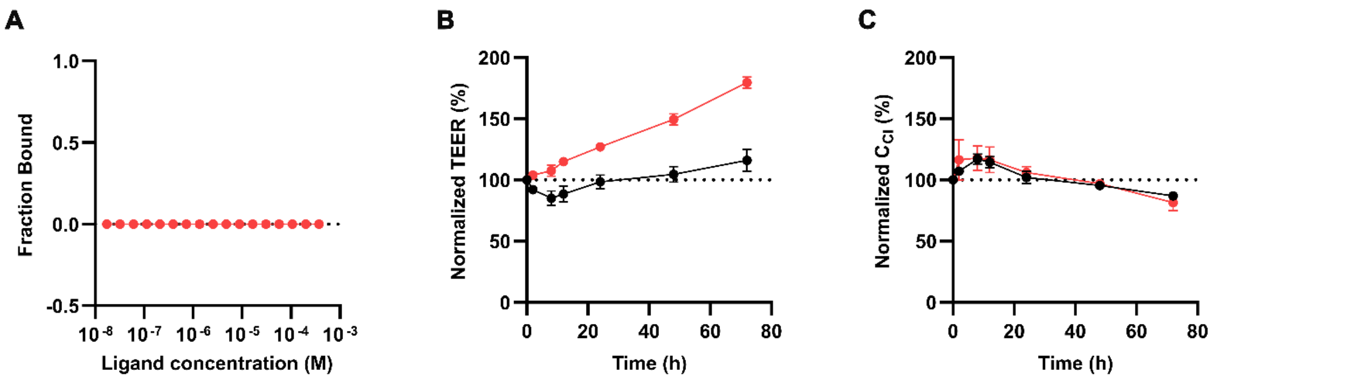

**Figure S6. Effect of SA14 exposure on binding and in vitro BBB properties. (A)** MST binding curves for SA14. No binding events were detected. (**B-C**) TEER (**B**) values and CCl (**C**), presented as percentages of baseline values, were measured in bEnd.3 monolayers treated with 250 μM SA14 (orange circles) for up to 72 h. The vehicle was used as a control condition (black circles).

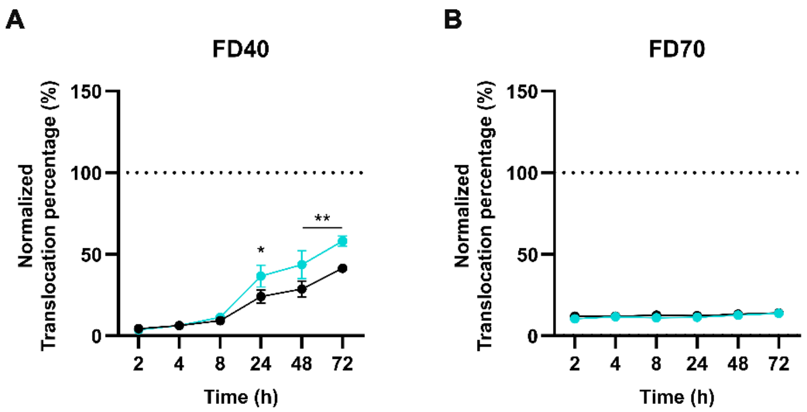

**Figure S7. ST9-mediated translocation of high-molecular-weight dextran across bEnd.3 cell layers.** (**A-B**) Normalized translocation ratio of FD40 (**A**) and FD70 (**B**) across bEnd.3 cell layers exposed to vehicle (Ctrl, black) or ST9 (250 μM, cyan circles). The translocation ratio, expressed as a percentage of translocation across a cell-free Transwell membrane, was calculated as the ratio between the dextran mass measured in the basolateral compartment at each sampling time point and the dextran mass initially loaded in the apical compartment at t = 0. Data are shown as means ± SEM (n = 3). *$p < 0.05$, **$p < 0.01$ vs Ctrl; two-way repeated-measures ANOVA/Tukey's tests.

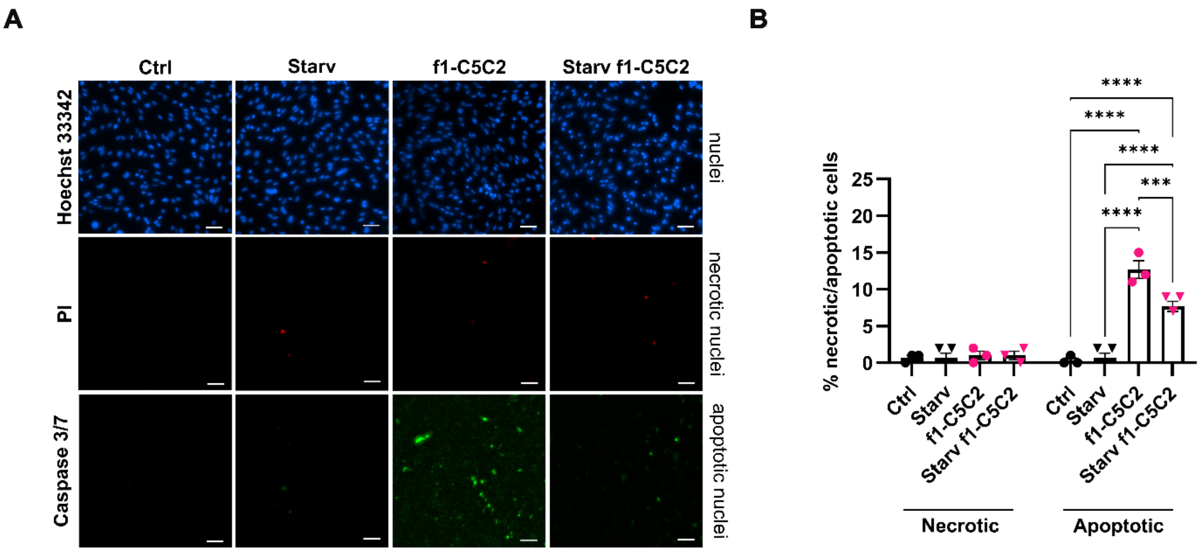

**Figure S8. Viability test upon exposure of bEnd.3 cells to the f1-C5C2 peptide and starvation.** (**A**) Representative fluorescence images at t = 4 h for bEnd.3 cells, cells were treated with vehicle (Ctrl, black) or 25 µM f1-C5C2 (pink) under non-starved conditions and compared with the corresponding starvation conditions (starvation alone: black; 25 ˜M f1-C5C2 under starvation: pink). Cells are stained with Hoechst 33342 (nuclear staining), Propidium iodide (PI, necrotic cells) and Caspase 3/7 (apoptotic cells). Scale bar, 50 µm. (**B**) Quantification of the amount of necrotic and apoptotic cells over total cells (%). Data are means ± SEM (n = 3 independent experiments; five-seven fields per replicate). ***$p < 0.001$, ****$p < 0.0001$; two-way repeated-measures ANOVA/Tukey's tests *versus* control.

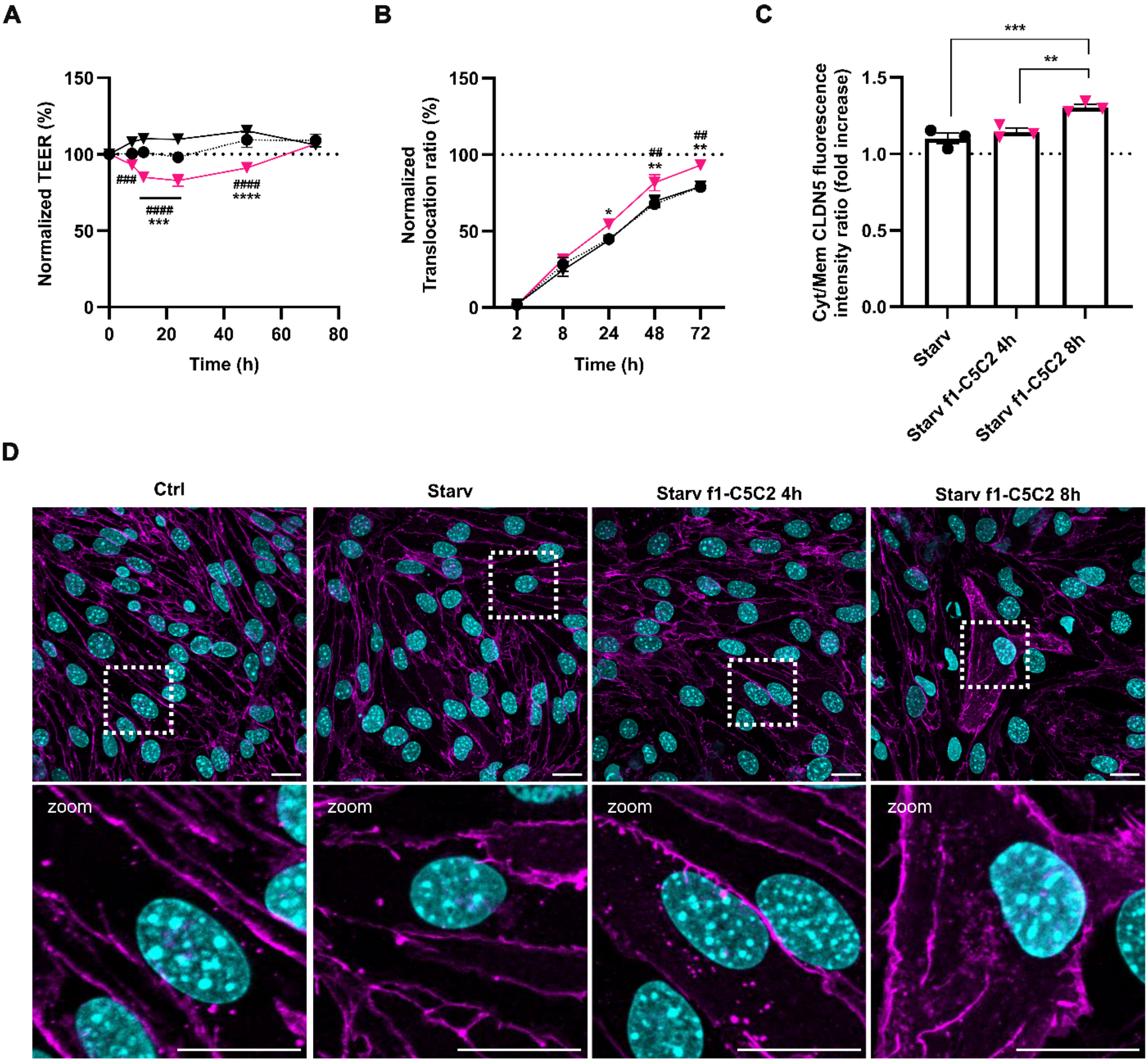

**Figure S9. Autophagy induction by starvation improves f1-C5C2-dependent BBB recovery and reduces CLDN5 internalization**. (**A-B**) TEER values (**A**) and normalized FD4 translocation ratio (**B**) were measured in bEnd.3 monolayers to evaluate the effect of starvation. Specifically, cells were treated with vehicle (Starv, black inverted triangles) or 25 µM f1-C5C2 (pink inverted triangles) under starvation and compared with the corresponding vehicle (Ctrl; black circles) under non-starved conditions for up to 72 h. Data are mean ± SEM (n = 3). *: Starv f1-C5C2 vs Ctrl; #: Starv f1-C5C2 *vs* Starv. *p < 0.05, **p < 0.01, ***p < 0.001, ****p < 0.0001 by two-way repeated-measures ANOVA with Tukey's multiple-comparisons test. (**C**) Quantification of CLDN5 intracellular localization based on image analysis. For each time point (4 and 8 hours), the cytoplasmic-to-plasma membrane CLDN5 fluorescence intensity ratio was calculated per cell and normalized to the ratio measured in matched controls. Data are means ± SEM (n = 3 independent experiments in triplicate; five fields per replicate). *p < 0.05, **p < 0.01, ordinary one-way ANOVA/Dunnett's test. (**D**) Representative confocal images of CLDN5 immunostaining in bEnd.3 cells treated with vehicle (Ctrl) or Starv and Starv f1-C5C2 conditions or with starvation induced, for 4 and 8 h. The boxed areas are shown at higher magnification under each image. Scale bar, 25 µm.

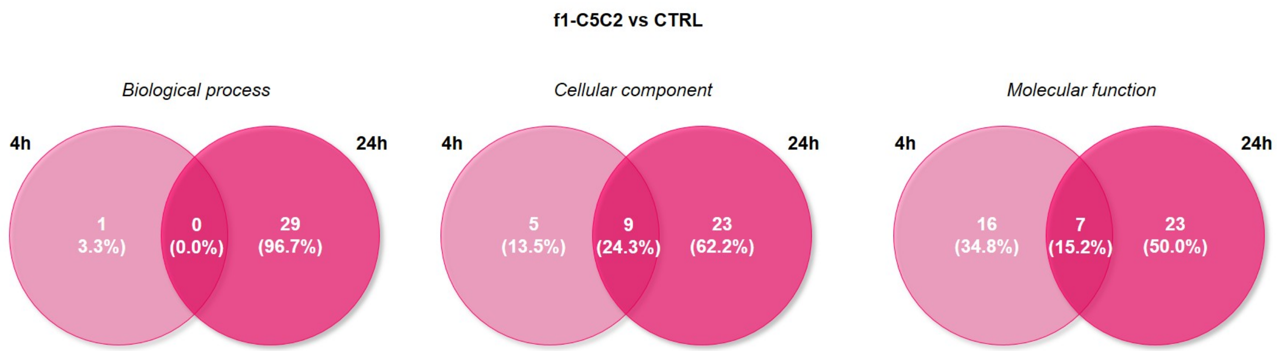

**Figure S8. Comparison of the effect of f1-C5C2 across different exposure times.** Venn diagrams of enriched pathways for biological process, cellular component, and molecular function after 4 h (light pink) and 24 h (dark pink) of exposure to f1-C5C2. The number of enriched pathways is higher after 24 h, indicating greater variations in protein abundance than in controls.

### Supplementary Tables

**Table S1. Hydrogen bonds formed by CLDN5-peptide complexes.** Persistence time of HBs formed between CLDN5 and ST9 ($d_1$-$d_9$) or SA14 ($d_{10-11}$). Only interactions with persistency > 20% of the total simulated time are reported. The results and associated errors are indicated as mean and standard deviation over three independent 500-ns-long replicas.

| *Distance* | *Complex* | *Residues (Atoms) involved* | *Persistency (%)* |
|---|---|---|---|
| $d_1$ | CLDN5-ST9 | Cys64($H_{N^*}$) – Ser6(O) | 97.7 $\pm$ 2.8 |
| $d_2$ | CLDN5-ST9 | Cys64(O) – Phe5($H_N$) | 91.0 $\pm$ 7.0 |
| $d_3$ | CLDN5-ST9 | Met62(O) – Ala7($H_N$) | 92.3 $\pm$ 8.1 |
| $d_4$ | CLDN5-ST9 | Met62($H_N$) – Ala7(O) | 80.2 $\pm$ 23.2 |
| $d_5$ | CLDN5-ST9 | Asp68($O\delta_1$) – Ser1($O\gamma$) | 24.1 $\pm$ 3.6 |
| $d_6$ | CLDN5-ST9 | Asp68($O\delta_2$) – Ser1($O\gamma$) | 23.0 $\pm$ 3.4 |
| $d_7$ | CLDN5-ST9 | Val66($H_N$) – Thr3(O) | 31.1 $\pm$ 3.2 |
| $d_8$ | CLDN5-ST9 | Gln63($O\varepsilon$) – Ser6($H\gamma$) | 29.0 $\pm$ 7.7 |
| $d_9$ | CLDN5-ST9 | Gly60(O) – Thr9($H_N$) | 72.3 $\pm$ 30.7 |
| $d_{10}$ | CLDN5-SA14 | Ala43(O) – Ala2($H_N$) | 22.5 $\pm$ 31.76 |
| $d_{11}$ | CLDN5-SA14 | Val56(O) – Trp3($H_N$) | 22.1 $\pm$ 31.2 |

**$H_N$: hydrogen atom bound to the backbone nitrogen*
